# TRANSITION OF PODOSOMES INTO ZIPPER-LIKE STRUCTURES IN MACROPHAGE-DERIVED MULTINUCLEATED GIANT CELLS

**DOI:** 10.1101/2020.02.26.966945

**Authors:** Arnat Balabiyev, Nataly P. Podolnikova, Aibek Mursalimov, David Lowry, Jason M. Newbern, Robert W. Roberson, Tatiana P. Ugarova

## Abstract

Macrophage fusion resulting in the formation of multinucleated giant cells (MGCs) is a multistage process that requires many adhesion-dependent steps and involves the rearrangement of the actin cytoskeleton. The diversity of actin-based structures and their role in macrophage fusion is poorly understood. In this study, we revealed hitherto unrecognized actin-based zipper-like structures (ZLSs) that arise between MGCs formed on the surface of implanted biomaterials. We established an *in vitro* model for the induction of these structures in mouse macrophages undergoing IL-4– mediated fusion. Using this model, we show that over time MGCs develop cell-cell contacts containing ZLSs. Live-cell imaging using macrophages isolated from mRFP- or GFP-Lifeact mice demonstrated that ZLSs are dynamic formations undergoing continuous assembly and disassembly and that podosomes are precursors of these structures. Immunostaining experiments showed that vinculin, talin, integrin α_M_β_2_, and other components of podosomes are present in ZLSs. Macrophages deficient in WASp or Cdc42, two key molecules involved in actin core organization in podosomes, as well as cells treated with the inhibitors of the Arp2/3 complex failed to form ZLSs. Furthermore, E-cadherin and nectin-2 were found between adjoining membranes, suggesting that the transition of podosomes into ZLSs is induced by bridging plasma membranes by junctional proteins.

## INTRODUCTION

Cell-cell fusion is a fundamental property of multicellular organisms and occurs in many physiological processes, such as fertilization, bone remodeling, skeletal muscle and placenta formation, and stem cell differentiation (Chen *et al*., 2007; Aguilar *et al*., 2013) (Podbilewicz, 2014). In addition, cellular fusion has been observed in numerous pathological conditions. In particular, the homotypic fusion of macrophages, leading to the formation of multinucleated giant cells (MGCs), occurs in tissues affected by chronic inflammation, including infectious and non-infections granulomas (Helming and Gordon, 2007b). Furthermore, MGCs are a prominent component of the foreign body reaction of the host to implanted biomaterials, and their accumulation at the tissue-material interface may persist throughout the lifetime of the implant (Anderson *et al*., 2008). MGCs adherent to biomaterials is known to produce potent cellular products that have been proposed to degrade the biomaterial, eventually leading to device failure (Zhao *et al*., 1991; Anderson *et al*., 2008) (Sheikh and Nash, 1996). MGCs are formed from blood monocytes recruited from the circulation to implant surface, where they differentiate into macrophages that undergo fusion. The T-helper 2 cytokine interleukin-4 (IL-4) participates in macrophage fusion *in vivo* (Kao *et al*., 1995) and is broadly used to study monocyte/macrophage fusion in cell cultures (McInnes and Rennick, 1988; McNally and Anderson, 1995) (Moreno *et al*., 2007) (Skokos *et al*., 2011), (Milde *et al*., 2015)

Cellular fusion is a multistage process, which starts with the cytokine induction of intracellular signaling that programs cells into a fusion-competent state. Adhesion of fusion-competent cells to a permissive substrate, cytoskeletal rearrangements, cell motility, and cell-cell interactions are all important determinants of macrophage fusion (Helming and Gordon, 2009). Most, if not all, of the steps involved in macrophage fusion, appear to rely on the actin cytoskeleton. It has long been known that cytochalasins B and D that prevent actin polymerization also inhibit macrophage fusion (DeFife *et al*., 1999) (Faust *et al*., 2019). The polymerization of actin filaments is known to be involved in the formation of diverse cellular protrusions, including lamellipodia, filopodia, and podosomes [for review see Ridley (2011) and Svitkina (2013)]. However, the precise targets of actin-disrupting agents that inhibit macrophage fusion have not been identified. Recently, we have characterized the early steps of IL-4–mediated macrophage fusion and showed that an actin-based protrusion at the leading edge initiates macrophage fusion (Faust *et al*., 2019). Furthermore, we have found that fusion-competent protrusions form at the sites enriched in podosomes.

Podosomes are dot-shaped adhesion complexes formed at cell-matrix contact sites that have been identified in many cell types. Podosomes are especially prominent in cells of the monocytic lineage, including macrophages and dendritic cells, where they are associated with cell adhesion, migration, and matrix degradation (for review see (Linder *et al*., 2011; Murphy and Courtneidge, 2011). Structurally, podosomes consist of a core of Arp2/3-mediated branched actin filaments and actin-regulatory proteins, including WASp and cortactin. The actin core is surrounded by a ring of cytoskeletal adaptor proteins, such as talin, vinculin, and paxillin. Moreover, integrin receptors have been localized in the ring (Zambonin-Zallone *et al*., 1989; Pfaff and Jurdic, 2001), with integrin α_M_β_2_ being the predominant integrin in macrophages (van den Dries *et al*., 2013b) and dendritic cells (Burns *et al*., 2004). The core and ring are linked by a myosin-IIA–containing network of unbranched actin filaments (Luxenburg *et al*., 2007; Akisaka *et al*., 2008; van den Dries *et al*., 2013b). An additional subset of unbranched actin filaments connects individual podosomes into groups (Bhuwania *et al*., 2012). Furthermore, a cap at the tip of the actin core consists of formins, fascin, and other proteins (Mersich *et al*., 2010; Van Audenhove *et al*., 2015) (Panzer *et al*., 2016), (Bhuwania *et al*., 2012). Podosomes are dynamic structures that give rise to different morphologies in diverse cells [for review see (Linder *et al*., 2011)]. In v-Src–transformed fibroblasts, podosomes form rosettes in cell extensions. In macrophage-derived osteoclasts, initial clusters of podosomes reorganize into rings, which then fuse and finally stabilize as a continuous sealing belt. Podosomes in macrophages are especially dynamic and undergo continuous turnover. Studies on the reorganization of podosomes in mononuclear mouse and human macrophages under nonfusogenic conditions have shown that individual podosomes can fuse with each other into larger structures and then disassemble into smaller clusters (Evans *et al*., 2003; Kopp *et al*., 2006; Poincloux *et al*., 2006). However, little is known about the fate of podosomes in MGCs as these giant cells undergo maturation *in vitro* and *in vivo*.

In this study, we revealed previously unknown actin-based zipper-like structures (ZLSs) that form between MGCs undergoing fusion on the surface of biomaterials implanted into mice. We established an *in vitro* model for the induction of these structures in MGCs formed by the IL-4– mediated fusion of mouse macrophages. Using this model, we found that podosomes were the precursors of ZLSs. Moreover, in addition to actin, ZLSs contained other proteins typically found in podosomes. The transition of podosomes into ZLSs appeared to be induced upon the bridging of two plasma membranes by the junctional proteins E-cadherin and nectin-2. Thus, a novel actin-based structure was identified in macrophages undergoing fusion on implanted biomaterials, which may provide a potential target for blocking MGC formation.

## RESULTS

### Zipper-like actin structures formed in MGCs following biomaterial implantation

While studying the foreign body reaction to implanted biomaterials in mice, we observed heretofore unrecognized actin-based structures formed at the contact sites between MGCs. In these experiments, polychlorotrifluoroethylene (PCTFE) sections were implanted into the peritoneal cavity of C57BL/6J mice, and the formation of MGCs was monitored after 3, 7, and 14 days by labeling recovered explants with Alexa Fluor 568-conjugated phalloidin and DAPI. Analyses of the samples revealed a progressive accumulation of MGCs on the biomaterial surface (Figure 1A). Furthermore, many MGC-MGC contact sites in the samples retrieved after 7 or 14 days of implantation contained areas with a seemingly symmetric and periodic actin distribution that visually resembled a zipper (Figures 1B). Based on this appearance, we hereafter call this actin pattern the zipper-like structure (ZLS). The ZLSs were first observed on day 7 as assessed by the measurements of the total ZLS length per high-power field (0.15 mm^2^), and their numbers increased by day 14 (Figure 1C). The majority of ZLSs were observed between MGCs, although they were infrequently seen at the contact sites of MGCs with mononuclear macrophages.

**Figure 1.**
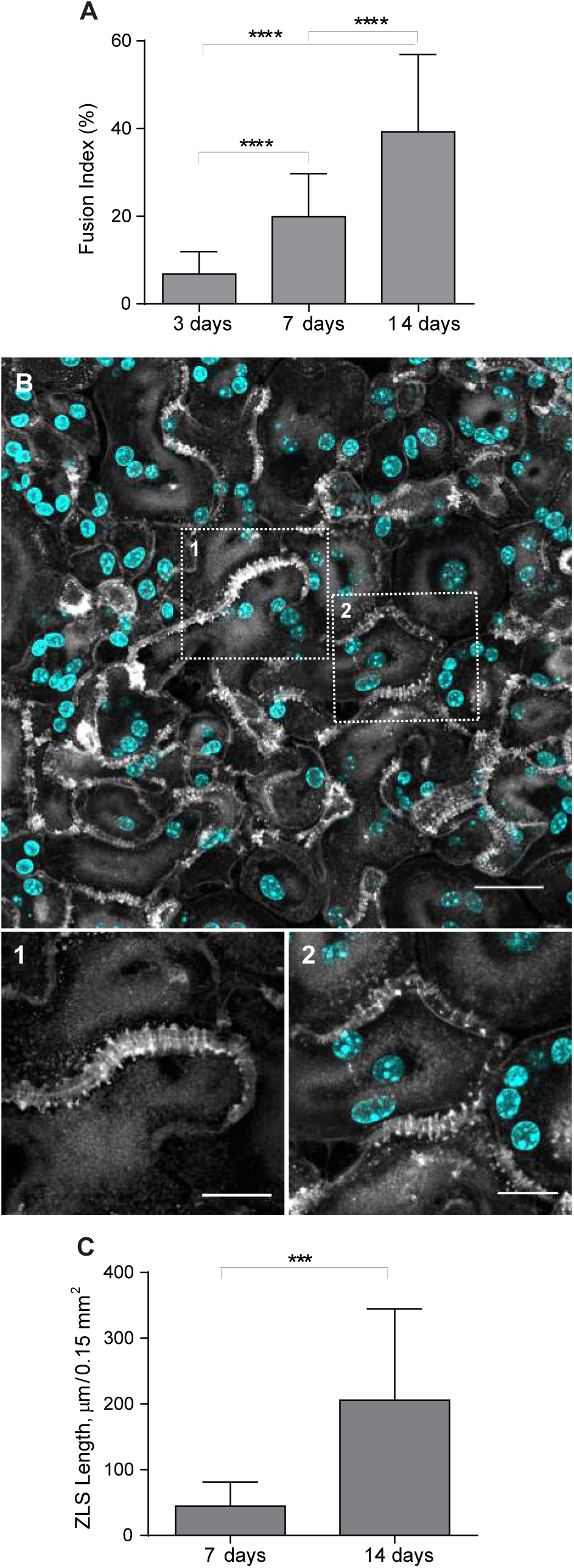
Formation of zipper-like structures (ZLSs) in MGCs following biomaterial implantation. **(A)** Sections of the PCTFE implanted into the peritoneal cavity of mice were recovered 3, 7, and 14 days after surgery. Explants were fixed and labeled with Alexa Fluor 546-conjugated phalloidin (white) and DAPI (teal). Macrophage fusion was assessed as a fusion index, which determines the percent of cells with three or more nuclei. Results shown are mean ± S.D. of three independent experiments. Three-to-five random 20× fields were used per sample to count nuclei. \*\*\*\**p* < .0001. **(B)** Representative images of MGCs formed on the surfaces of the implants recovered at day 14 post-surgery. The scale bar is 20 µm. *Bottom panels*, high magnification views of the boxed areas in A. The scale bars are 10 µm and 15 µm, respectively. **(C)** The time-dependent formation of ZLSs on the PCTFE sections explanted at days 7 and 14 post-surgery. The formation of ZLSs was assessed as the total length of ZLSs per high-power field (0.15 mm^2^), and the determination was made using ImageJ. Results shown are mean ± SD of three independent experiments. Three-to-five random 20× fields were used per sample to count nuclei. \*\*\**p* < .001.

### Formation of ZLSs *in vitro*

To investigate the mechanism of ZLS formation, we established an *in vitro* system that allowed us to generate ZLSs reproducibly. Since PTFE plastic is not amenable to most imaging techniques, we took advantage of recently developed optical-quality glass surfaces prepared by adsorption of long-chain hydrocarbons such as paraffin that promote high level of macrophage fusion (Faust *et al*, 2017; Faust *et al*., 2018). In this series of experiments, rather than macrophage cell lines, we used primary macrophages isolated from inflamed mouse peritonea (Helming and Gordon, 2007a; Podolnikova *et al*., 2016; Faust *et al*., 2019) to avoid the robust proliferation observed in the cultures of macrophage cell lines. Using phase-contrast video microscopy, we determined that the kinetics of IL-4–induced macrophage fusion on paraffin-adsorbed glass (P-surface) were similar to that on PCTFE (Figure 2A and B). The cell fusion began 8–9 h on both surfaces after the addition of IL-4, and the maximum number of fusion events occurred after 16–20 h and then gradually declined. Furthermore, as determined by the measurements of the fusion index, the period between hours 9 and 24 was the most active period of fusion on both surfaces (Figure 2C). Based on this similarity, we used P-surfaces in subsequent experiments. Since we were unable to detect ZLSs after 48 h, the duration of incubation was extended to 5 days. Although the size of MGCs did not significantly change after 2 days (Supplemental Figure 1), they acquired a more round morphology by day 4 relative to the irregularly shaped MGCs formed at earlier times (Figure 2D). In addition, contacts between apposing MGCs became more frequent, with many regions displaying ZLSs (Figure 2D and F, F’). ZLS formation appeared to have correlated with the fusion index inasmuch as only the limited number of ZLSs was detected on poorly fusogenic acid-cleaned glasses (Fig. 2E, G and H). The total length of ZLSs at day 5 ranged between a minimum of ∼50 µm and a maximum of ∼1460 µm/high power field (Figure 2I), and the average length of individual ZLSs was 35 ± 25 µm and varied between 6 µm and ∼150 µm (Figure 2J). Furthermore, increasing the seeding cell density resulted in an increase in the total ZLS length (Figure 2K), suggesting that contact between cells is essential for ZLS formation. After 5 days, the MGCs and mononuclear macrophages attached to P-surface remained viable as determined by the trypan blue exclusion test (Supplemental Table 1). Based on the examination of confocal images (Figures 1B and 2, D and F), the pattern of actin distribution within ZLSs formed *in vitro* and *in vivo* appeared to be indistinguishable. In subsequent experiments, the 5-day time point was chosen to allow for sufficient ZLS formation.

**Figure 2.**
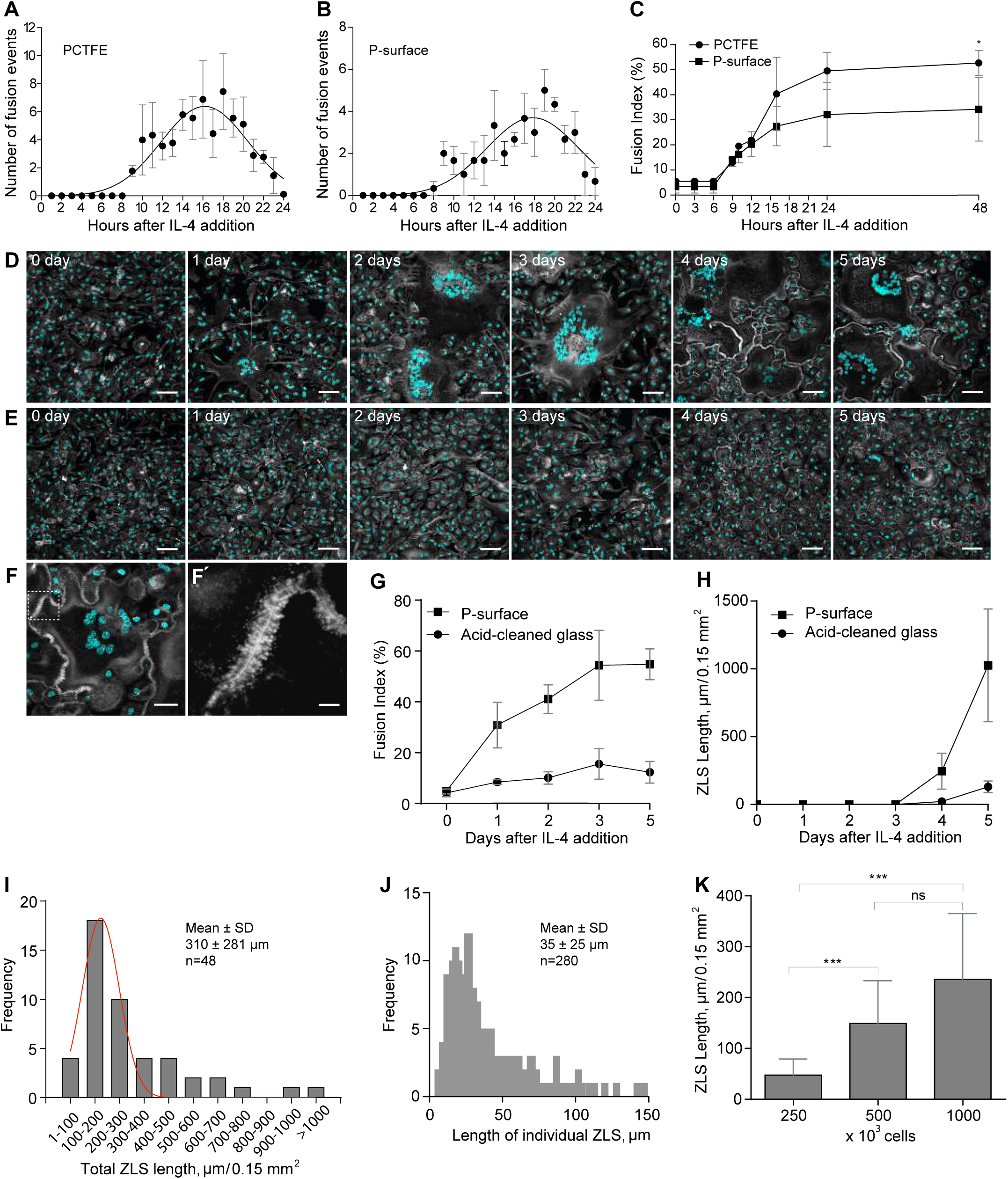
Kinetics of macrophage fusion and ZLS formation *in vitro*. **(A, B)** Macrophages were isolated from the mouse peritoneum 3 days after TG injection and plated on PCTFE sections (20 × 20 mm) or paraffin-coated cover glasses (P-surface) at 5000 cells/mm^2^ in DMEM/F12 supplemented with 10% FBS, and fusion was induced by the addition of IL-4 (10 ng/ml). The number of fusion events at different times was determined using phase-contrast video microscopy. Results shown are mean ± SD of three-four independent experiments. **(C)** The fusion indices of macrophages plated on PCTFE or P-surface determined at different time points. Results shown are mean ± SD of three independent experiments. Three-to-five random 20× fields were used per sample to count nuclei (total ∼6000 nuclei). **(D, E)** TG-elicited peritoneal macrophages were seeded at 5000 cells/mm^2^ on paraffin-coated (**D**) or acid-cleaned (**E**) cover glasses, and fusion was induced by the addition of IL-4 (10 ng/ml). Representative images of fusing macrophages taken at different time points are shown. The scale bars are 50 μm. **(F)** A representative image of an MGC formed in the 5-day culture of fusing macrophages on the P-surface. The scale bar is 50 μm. **(F’)** High magnification view of a boxed area in **F**. The scale bar is 5 µm. **(G)** The time-dependent fusion of IL-4–induced macrophages on the P-surface or acid-cleaned glass. Results shown are mean ± SD of three independent experiments. **(H)** The total lengths of ZLSs in MGCs formed on the P-surface or acid-cleaned glass for 5 days. Results shown are mean ± SD of three independent experiments. Three-to-five random 20× fields were used per sample to determine the ZLS length by Image J. **(I)** The frequency distribution of the total ZLS lengths/high power field in MGCs formed in the 5-day culture (n = 48). **(J)** The frequency distribution of individual ZLS lengths (n = 280). **(K)** The total lengths of ZLSs formed in the 5-day cultures of macrophages plated at different densities. Results shown are mean ± SD of three independent experiments. Three-to-five random 20× fields were used per sample to determine the length. \*\*\**p* < .001; ns, nonsignificant.

### The three-dimensional pattern of the actin distribution in ZLS

To examine whether ZLSs had a specific pattern, we determined their dimensional parameters using samples from 5-day MGC cultures labeled with Alexa Fluor 568-conjugated phalloidin. The periodicity of the actin distribution in ZLSs was determined from the x-y planes (Figure 3A), and the height and width from the scans of fluorescence intensity of the x-z sections (Figure 3, B and C). In a ZLS cross-section, actin was organized into large and small globules that formed two closely spaced “humps” originating from each MGC (Figure 3C). The average maximum height of the humps was 2.9 ± 0.5 µm (n = 64; 40 cells), and the average width was 4.8 ± 0.9 µm (n = 196; 30 cells). The distribution of the height and width values of the actin humps is shown in Supplemental Figure 2. The humps were closely abutting at the site of cell-cell contact. (Figure 3C). The average height of the region of close apposition was 1.2 ± 0.3 µm (n = 40; 20 cells). The average periodicity of the main actin foci seen in ZLSs was 2.1 ± 0.4 µm (n = 60; 30 cells) (Figure 3B, arrowheads and Figure 3F). By fitting the diameter value distribution of the bottommost region of the large globules with a bimodal Gaussian formula, two populations were identified (Figure 3G) with average diameters of 1.2 ± 0.2 µm and 2.0 ± 0.3 µm (n = 100). Another feature observed in the x-y plane was the areas of actin organization that appeared as closely spaced small globules lying along the plasma membrane of two apposing MGCs (Figure 3B, arrows). The images acquired by structured illumination microscopy (SIM) showed additional details of this area (Figure 3, D and E). The space between the plasma membranes was clearly seen with small actin globules (0.24 ± 0.06 µm in diameter) positioned at seemingly regular intervals (0.2 ± 0.1 µm) at the cytosolic face of the membrane (Figure 3H). In the x-y plane, it appeared that large foci were connected to small foci by thin filaments, although such filaments were not clearly seen in some locations. Together, these analyses suggest that the actin distribution pattern in ZLS is relatively uniform.

**Figure 3.**
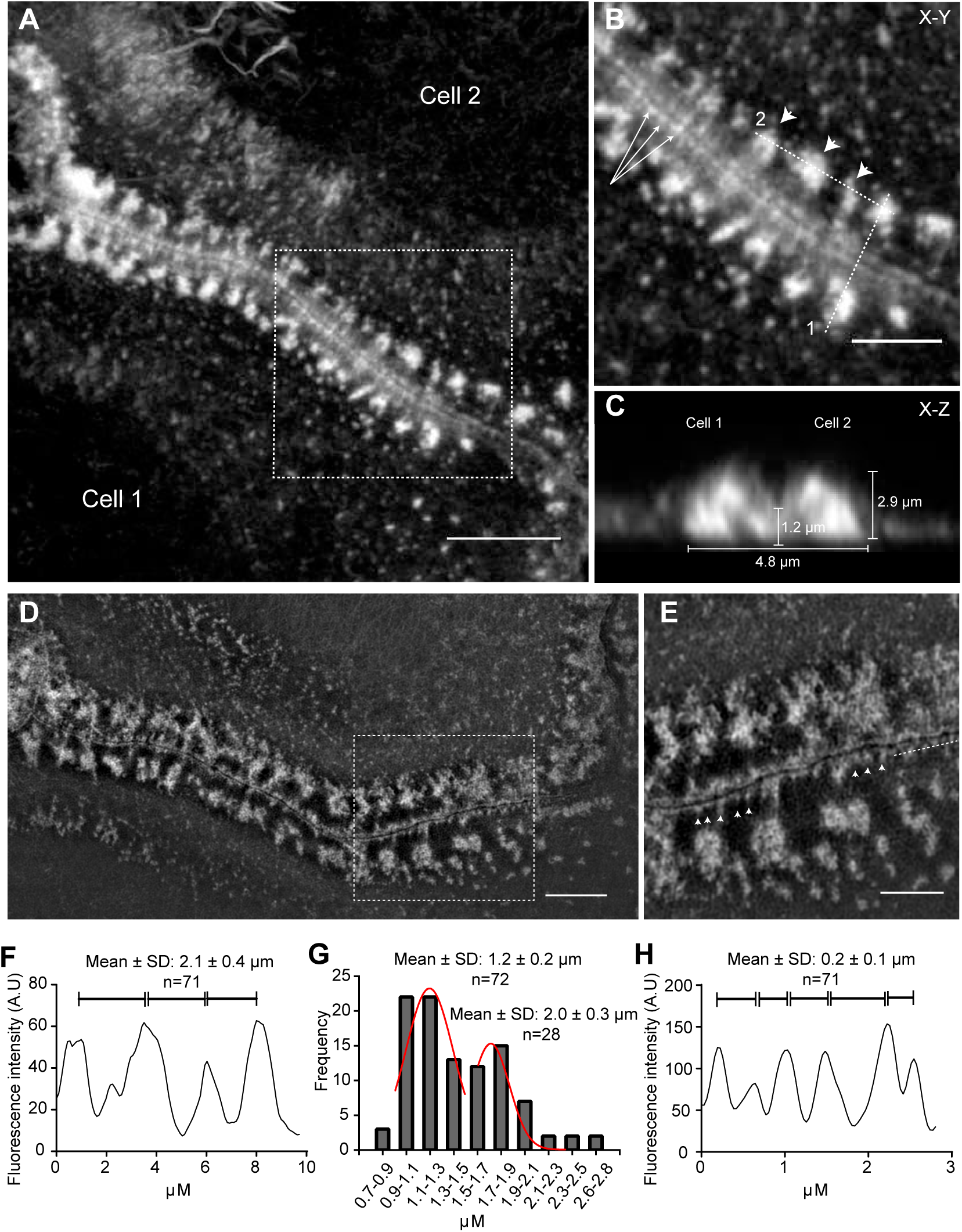
Three-dimensional pattern of actin distribution in ZLS. **(A)** MGCs formed in a 5-day culture were labeled with Alexa Fluor 568-conjugated phalloidin, and confocal images were used to analyze actin distribution in ZLSs. A representative deconvolved x-y confocal plane of a ZLS close to the substrate is shown. The scale bar is 5 μm. **(B)** High magnification image of the boxed area in A. The arrowheads point to the main actin foci, and arrows point to small actin globules adjacent to the plasma membranes. The scale bar is 2.5 μm. **(C)** The z-scan through line 1 in **B** demonstrates a cross-section through the ZLS and illustrates the localization of actin in two adjacent humps. Numbers indicate the averaged dimensional parameters of the ZLS. **(D)** Imaging of MGCs labeled with Alexa Fluor 568-conjugated phalloidin using SIM. The scale bar is 5 μm. **(E)** A high magnification image of the boxed area in **D**. Arrowheads indicate membrane-adjacent actin globules. The scale bar is 2.5 μm. **(F)** The scan of fluorescence intensity (in arbitrary units) along dotted line 2 indicated in B. **(G)** The size distribution of the large actin globules. **(H)** The scan of the fluorescence intensity (in arbitrary units) along a dotted line shown in E.

### ZLSs are transient actin-based arrangements formed from podosomes

Upon close examination of adjoining cells (Figures 4), we noticed that the regions of the plasma membranes just preceding the ZLSs (Figure 4, enlarged boxes 1 and 2, arrows) contained actin puncta that apparently represented podosomes, a characteristic feature of macrophages. Furthermore, individual podosomes were seen in the vicinity of these sites (Figure 4, enlarged boxes 1 and 2, *arrowheads*). Mononuclear macrophages that were often observed close to MGCs also contained numerous circumferential podosomes (Figure 4, *left bottom quadrant*). While mononuclear macrophages occasionally formed short ZLSs with MGCs (Figure 4, enlarged box 3), they did not form ZLSs with each other. The presence of podosomes in close proximity to the site of the MGC-MGC apposition suggested that podosomes might be precursors of ZLSs.

**Figure 4.**
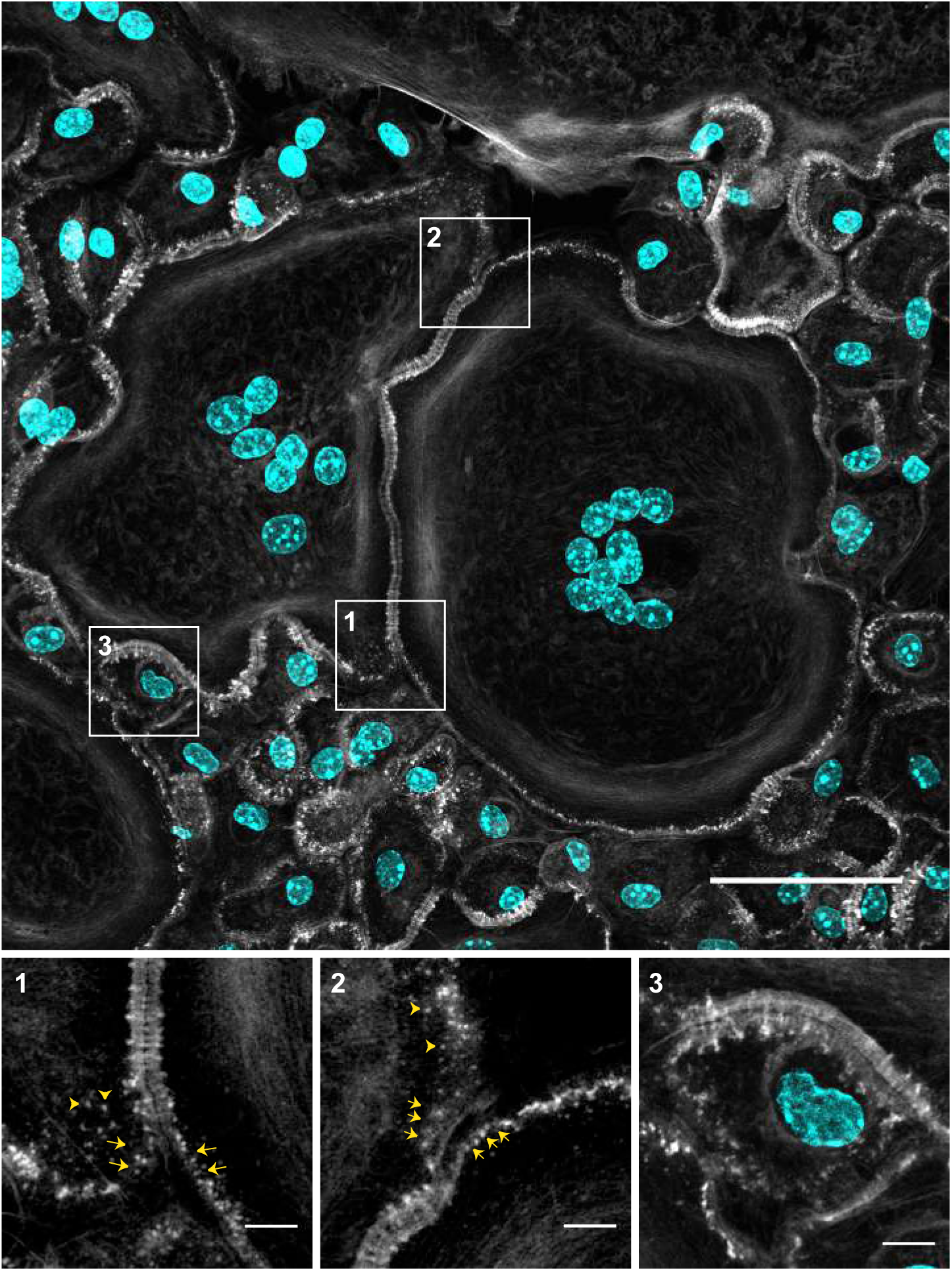
Podosomes as potential precursors of ZLSs. A representative image of MGCs formed in the 5-day culture of fusing macrophages. Cells were labeled with Alexa Fluor 488-conjugated phalloidin (white) and DAPI (teal). Many areas of contact between MGCs and MGC-mononuclear cells contain ZLSs, which are observed at the sites where two plasma membranes decorated with podosomes adjoin. The scale bar is 50 μm. High magnification images of the boxed areas **(1-3)** illustrate the possible formation of ZLSs from podosomes. The scale bars are 5 μm.

To examine whether ZLSs originate from podosomes, we performed live-cell imaging using mRFP- or GFP-LifeAct macrophages, which were fused by IL-4 induction for 5 days. Surprisingly, ZLSs were not as stable as appeared in the fixed specimens analyzed by immunofluorescence but were rather dynamic. The ZLSs formed from actin puncta remained in the zipper for some time and then disassembled into actin puncta. As shown in Figure 5A and Video 1, a nascent ZLS (00:00 min; yellow and white arrows show the top and bottom borders of the ZLS, respectively) was undergoing further organization and zipper elongation (2:00–06:00 min). The subsequent growth of this ZLS downward from a pool of adjacent podosomes was concomitant with its disassembly upstream (Figure 5A, 10:00–22:00 min). A cloud of actin puncta emerging from the disassembled ZLS remained near the plasma membrane. Another area of the ZLS assembly and disassembly is shown in Figure 5B and Video 2. The dismantlement of the ZLS, in this case, resulted in the separation of the two MGCs (Figure 5B; 35:00 min). The amount of time during which the membranes between different MGCs remained joined by ZLSs was 12.8 ± 3.5 min (n = 20). No correlation between the length of the ZLSs and time was found (Supplemental Figure 3).

**Figure 5.**
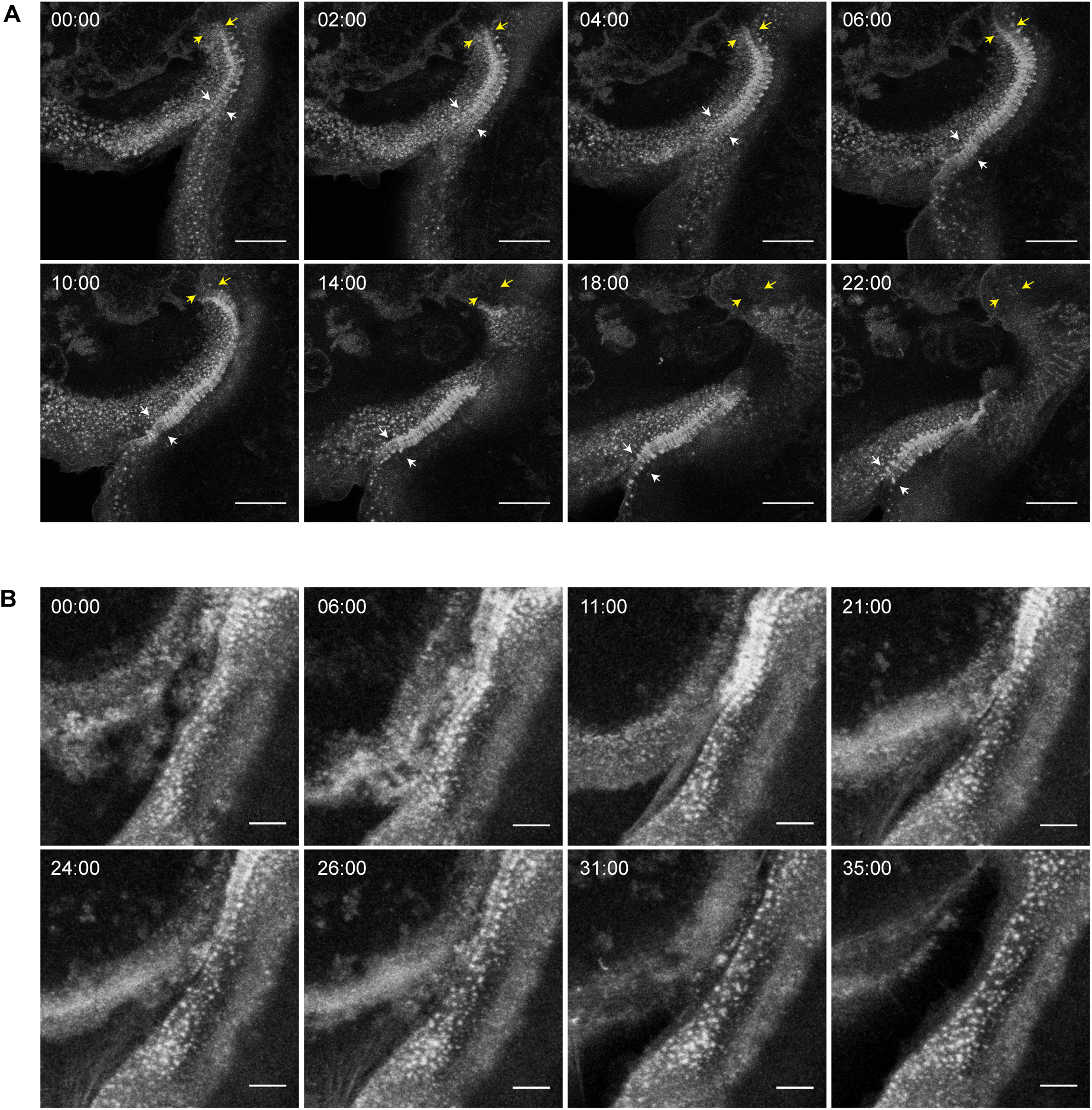
Live-cell imaging of ZLS formation. **(A)** Frames from a representative live-cell imaging experiment showing the formation of a ZLS between two MGCs formed in a 5-day culture of IL-4–induced macrophages isolated from mice expressing mRFP-LifeAct. Yellow and white arrows indicate the initial upper and bottom boundaries of the nascent ZLS, respectively (00:00 min). After the period of maturation and elongation (02:00-10:00 min), the upper part of the ZLS disassembles (10:00-22:00 min). The scale bar is 20 μm. See also Video 1. **(B)** Frames from a movie showing the formation and disassembly of another ZLS between MGCs derived from EGFP-LifeAct macrophages The scale bar is 5 μm. See also Video 2.

### ZLSs contain proteins found in podosomes

Further evidence for the podosome origin of ZLSs was obtained by immunofluorescence experiments using antibodies that recognize proteins usually found in podosomes. The defining feature of podosomes is the presence of the core of actin filaments (Linder *et al*., 2000; Kaverina *et al*., 2003) surrounded by adhesive plaque proteins, such as talin, vinculin, and integrins (Zambonin-Zallone *et al*., 1989) (Pfaff and Jurdic, 2001). As shown in Figure 6, vinculin, talin, paxillin, cortactin, and integrin α_M_β_2_ were detected in ZLSs. Furthermore, actin and podosome proteins were enriched on both sides of the plasma membranes. Among the podosome proteins, vinculin and talin localized in the humps, and both proteins encircled the large actin globules (Figure 6, A-F). No colocalization of vinculin or talin with actin was observed in the small globules (Figure 6, B and E). Paxillin was also found to surround the large globules (Figure 6, G and G’). Cortactin colocalized with actin in the large globules and was present in the small globules in close proximity to the midline between the two MGC membranes (Figure 6H and H’; arrow in the X-Z section). The area between the humps corresponding to the site of close apposition between the plasma membranes appeared to be void of podosome-associated proteins, except for α_M_β_2_ (Figure 6I and I’). The integrin α_M_β_2_ was present in the midline and also decorated the plasma membrane above the humps. Myosin II, which has been shown to localize in the area surrounding the actin core (Labernadie *et al*., 2010; van den Dries *et al*., 2013a) and to enrich in the regions of high podosome turnover (Kopp *et al*., 2006), was also found in ZLSs (Supplemental Figure 4A).

**Figure 6.**
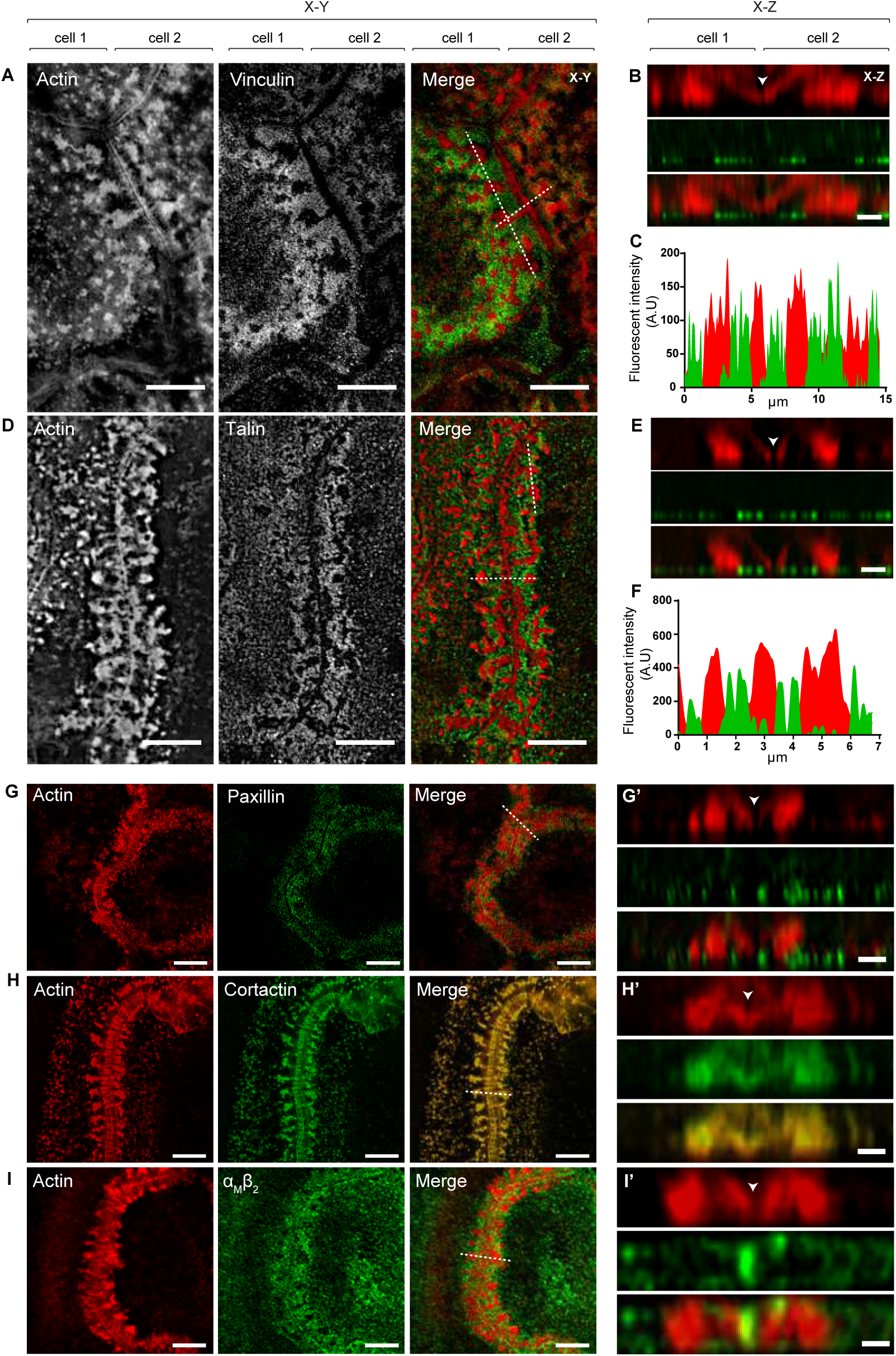
Localization of podosome-specific proteins in ZLSs. Representative SIM images of MGCs in the 5-day culture induced by the addition of IL-4 (10 ng/ml) and incubated with anti-vinculin **(A),** anti-talin **(D),** anti-paxillin **(G),** anti-cortactin **(H)** and anti-α_M_β_2_ **(I)** antibodies followed by corresponding secondary antibodies conjugated to Alexa Fluor 488. Actin was labeled with Alexa Fluor 568-conjugated phalloidin. Confocal x-y planes close to the substrate-attached cell surface show the distribution of actin and podosome-specific proteins. **(B, E, G’, H’, I’)** The z-scans across the horizontal dotted lines in the merged images shown in **A, D, G, H,** and **I** illustrate the distribution of actin and podosome-specific proteins in the humps. The midline, which defines the position of adjoining plasma membranes, is indicated by arrowheads. **(C, F)** Scans of fluorescence intensity of actin/vinculin and actin/talin, respectively, across the vertical dotted lines (x-y plane) shown in A and D. The scale bars are 5 μm in all x-y sections and 2 µm in all x-z sections.

It is known that the Arp2/3 complex and its activators Cdc42 and WASp nucleate the core of actin filaments in podosomes (Linder *et al*., 2000). We have previously demonstrated that deficiency of WASp or Cdc42, or inhibition of Arp2/3 strongly reduces macrophage fusion and podosome formation (Faust *et al*., 2019). To examine the importance of these actin regulators in ZLS formation, we isolated macrophages form WASP^-/-^ or myeloid-cell–specific Cdc42^-/-^ mice and tested their ability to assemble ZLSs. We observed that WASp- (Figure 7, A, B, E, and F) or Cdc42-deficient macrophages (Figure 7, C, G, and H) lost their capacities to form ZLSs in correlation with their reduced fusion capacities. In addition, the treatment of wild-type macrophages with wiskostatin, a specific inhibitor of WASp, decreased the fusion and total length of the ZLSs (Figure 7D and E, F). Next, we examined whether inhibition of Arp2/3 impaired the ability of wild-type macrophages to form ZLS. As shown in Figure 7I, the Arp 2/3-specific inhibitor CK-636 or CK-548 blocked ZLS formation in a dose-dependent manner. Together with live-cell video microscopy results, these data suggest that ZLSs originate from podosomes.

**Figure 7.**
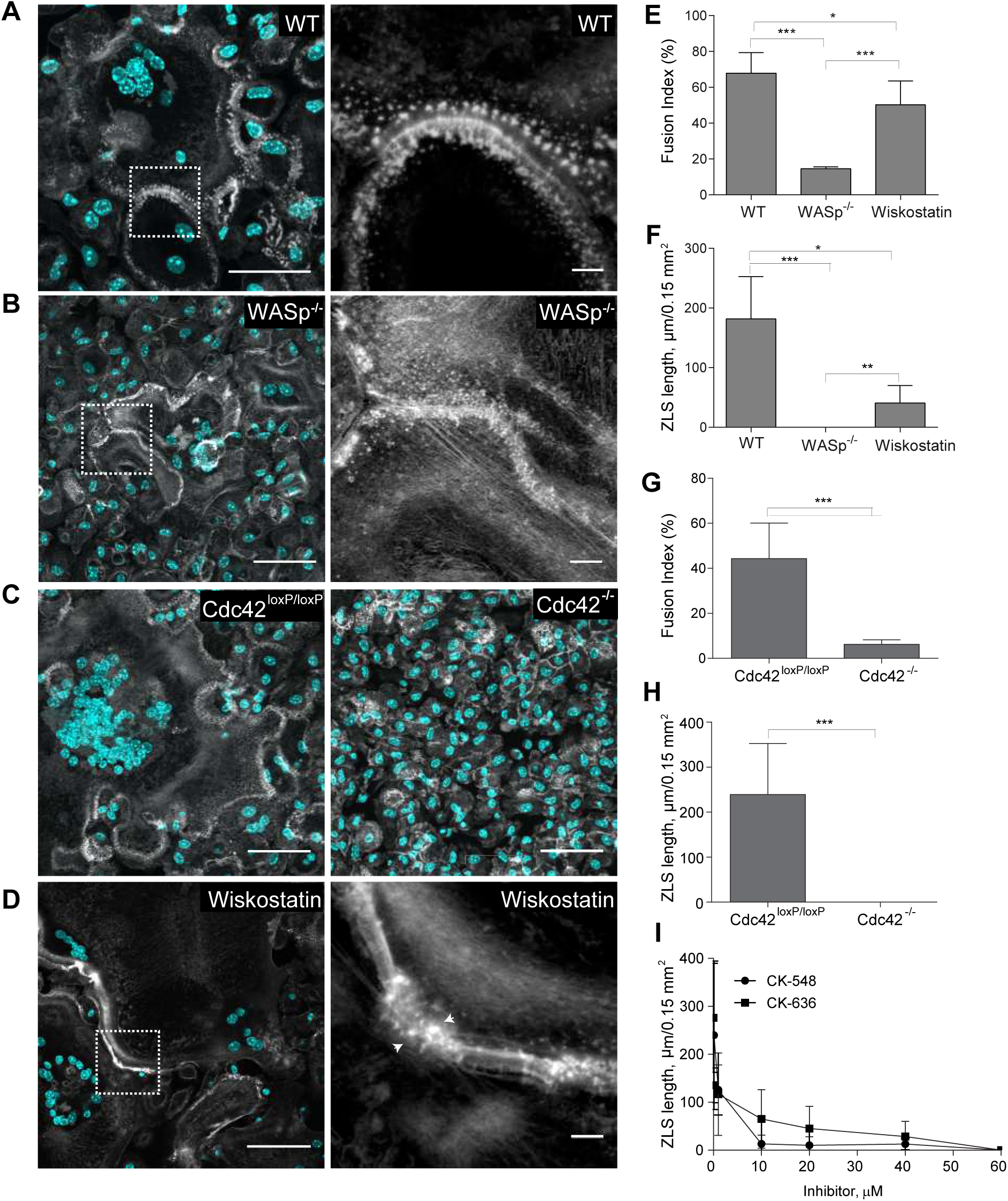
The role of organizers of actin polymerization in ZLS formation. Formation of ZLS in the 5-day cultures of wild-type **(A),** WASp-deficient, **(B)** or Cdc42-deficient **(C)** macrophages induced by the addition of IL-4 (10 ng/ml). The right panels in A and B show high magnification images of the boxed areas in the left panels. The scale bars are 50 μm and 5 μm for the original and high magnification images, respectively. **(D)** Effect of wiskostatin (1 µM) on the formation of ZLS. Wiskostatin was added simultaneously with IL-4. **(E-H)** Fusion index and the lengths of ZLSs in MGCs derived from WASp- or Cdc42-deficient macrophages or in macrophages treated with wiskostatin. Results shown are mean ± SD of three independent experiments. \**p* < .05, \*\**p* < .05, \*\*\**p* < .001 **(I)** Effect of the Arp2/3 inhibitor CK-548 or CK-636 on ZLS formation. The total lengths of ZLSs per high power field (0.15 mm^2^) were determined in MGCs formed in the 5-day cell culture.

### Analyses of the junctional proteins within ZLS

Analyses of actin distribution shown in Figure 3 revealed a narrow space between adjoined MGCs, suggesting a very close apposition between the plasma membranes within ZLSs. To examine the interface between the membranes at a higher resolution, we performed transmission electron microscopy (TEM) to examine MGCs formed in the 5-day culture. Analyses of the sections taken parallel the site of the cell attachment to substrate revealed segments of closely apposed plasma membranes with a spacing of 7.7 ± 1.3 nm (n= 30) (Figure 8A). Interestingly, the space between two membranes was not empty but filled with a material, which displayed striation in some areas (Figure 8A, *inset* in the left panel). To determine the nature of these electron-dense “ladders”, we performed confocal microscopy using antibodies against selected proteins, including E-cadherin, nectin-2, JAM-A, and connexin43, which are known to mediate homophilic cell-cell interactions (Figure 8, B-F). The presence of these proteins has previously been reported in activated macrophages (Liu *et al*., 2000; Eugenin *et al*., 2003; Pende *et al*., 2006; Moreno *et al*., 2007; Van den Bossche *et al*., 2009). We also labeled cells with the lipophilic membrane stain DiD to mark the midline, which separates the two halves of ZLSs (Figure 8B, *arrowhead in the right panel*). Labeling with a mAb against the ectodomain of E-cadherin revealed that this protein was present on ZLSs (Figure 8C). E-cadherin was detected in the midline (x-y plane) and at the apposition site between the humps (x-z section). E-cadherin was also present at the ventral sides of the humps but was mainly excluded from the bodies of the humps. Both nectin-2 (Figure 8D) and JAM-A (Figure 8E) were detected in ZLSs. However, only nectin-2 was found in both the midline and at the apposition site, whereas JAM-A appeared to have colocalized with actin in the humps. Labeling for afadin, a cytosolic adaptor protein which associates with nectin-2 (Takai *et al*., 2008), showed its colocalization with actin in the humps (Figure 8F). Connexin43 was not detected on ZLSs. Since the tight junction protein ZO-1 has also been found in macrophages, we have labeled ZLSs with anti-ZO-1 antibody; however, we were unable to detect this protein.

**Figure 8.**
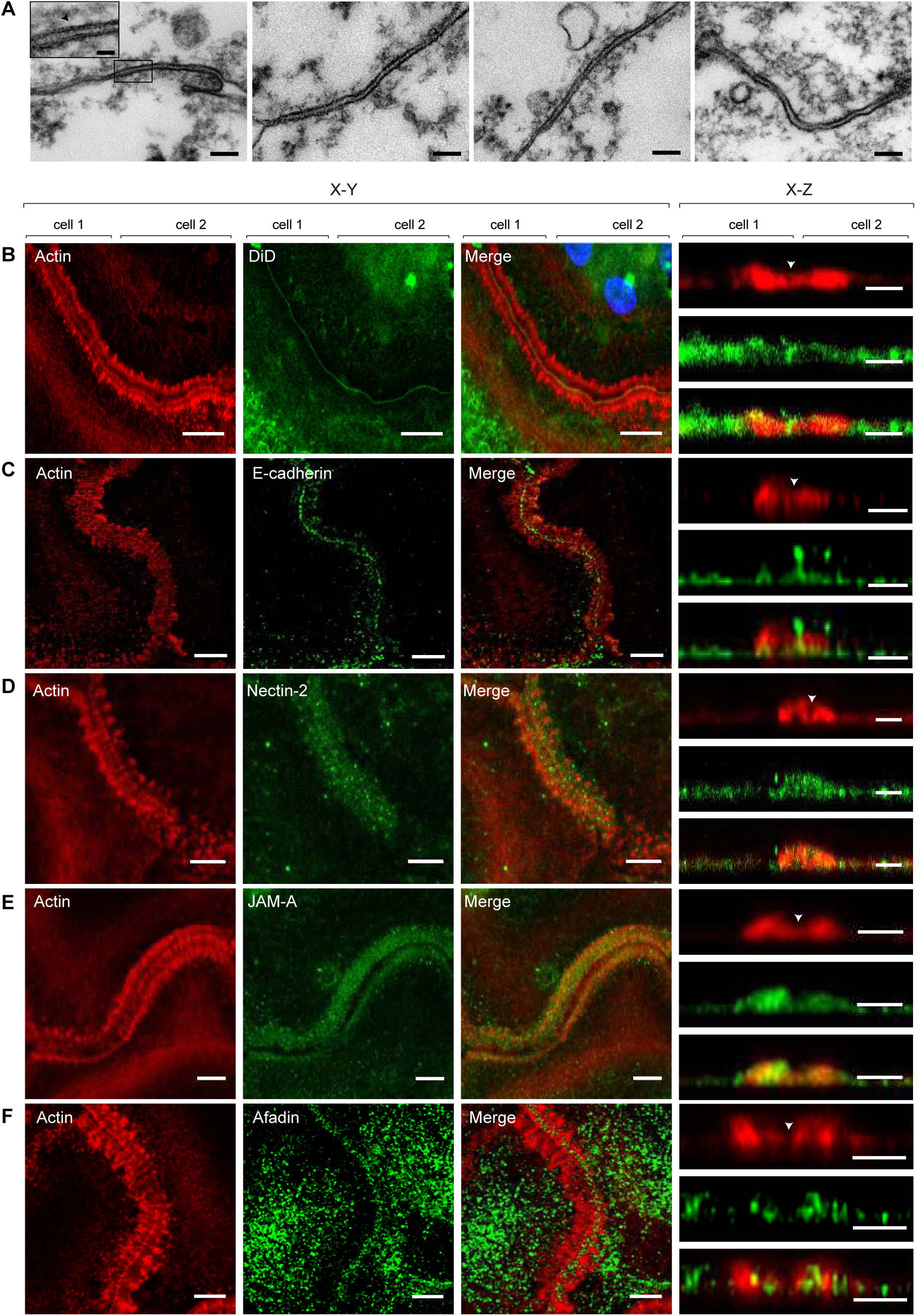
Analyses of junctional proteins within ZLS. **(A)** Ultrastructural details of the intercellular junctions formed in a 5-day MGC examined by TEM. Representative TEM micrographs of the sections prepared by cutting the specimen parallel to the substratum. The scale bar is 100 nm. *Inset* in the left panel is a higher magnification view of the boxed area. **(B-F)** MGCs in the 5-day culture were labeled with DiD **(B)** or incubated with anti-E-cadherin **(C),** anti-nectin-2 **(D),** JAM-A, **(E)** or anti-afadin **(F)** antibody followed by corresponding secondary antibodies conjugated to Alexa Fluor 488. Actin was labeled with Alexa Fluor 568-conjugated phalloidin. The x-y confocal planes close to the cell-substratum attachment side (200 nm; *left panels*) and the x-z sections of the z stack (*right panels*) are shown. The scale bars are 5 µm and 2.5 μm for the x-y and x-z planes, respectively.

Investigations of E-cadherin and nectin-2 expression during macrophage fusion showed that E-cadherin (Figure 9A) and nectin-2 mRNAs (Supplemental Figure 5) were barely expressed in freshly isolated macrophages. E-cadherin transcript was detected at hour 3 of incubation of macrophages in the presence of IL-4. A ∼10-fold increase was observed at hour 12, and this level did not change after 5 days (Figure 9B). Likewise, E-cadherin was poorly expressed on the surface of macrophages before their treatment with IL-4 but gradually increased after the addition of IL-4 (Figure 9, C and D).

**Figure 9.**
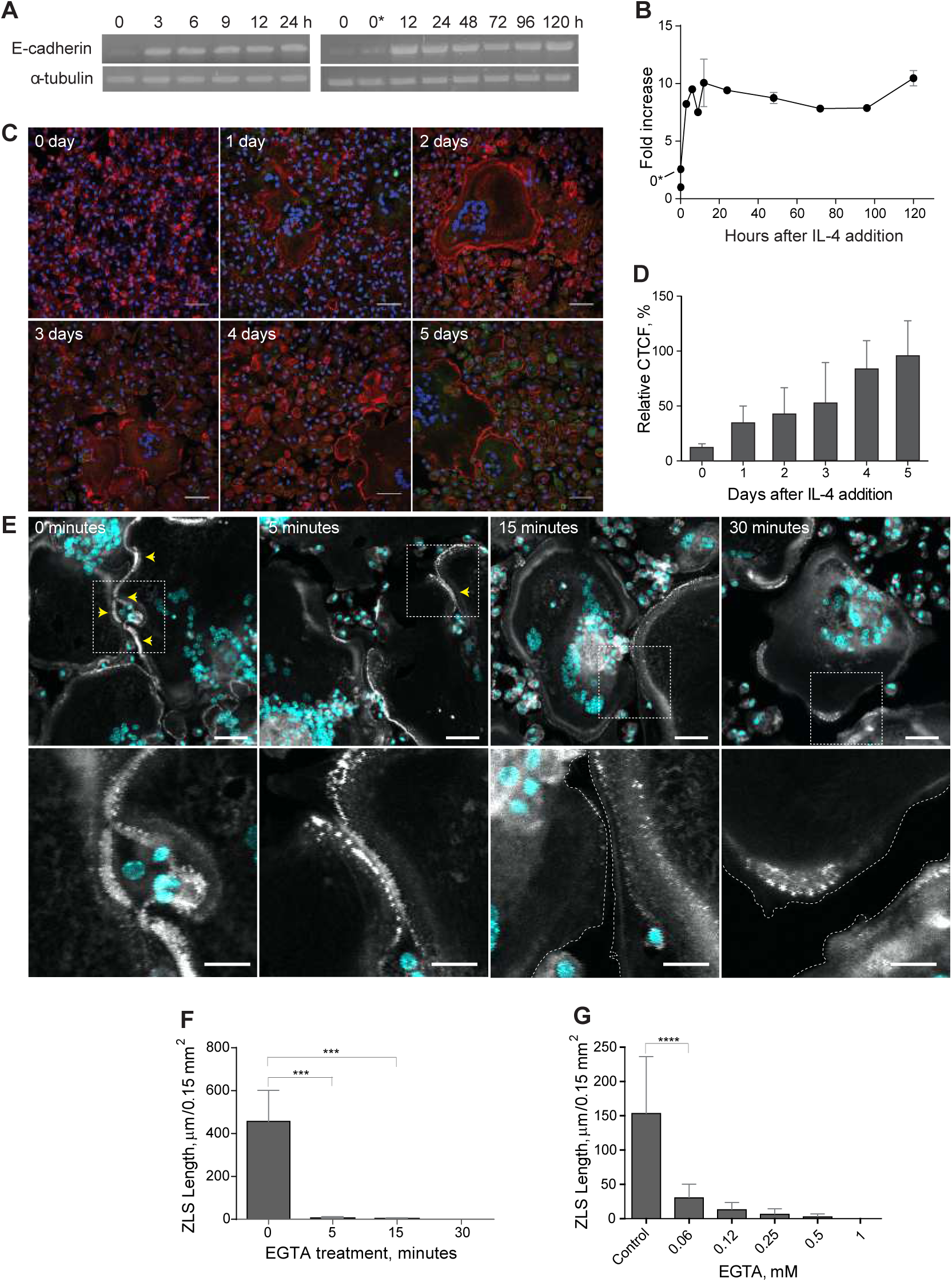
Analyses of E-cadherin expression in fusing macrophages and its role in ZLS formation. **(A, B)** A time course of E-cadherin mRNA expression in macrophages undergoing fusion in the presence of IL-4, as determined by RT-PCR. Signal intensities were normalized to that of α-tubulin mRNA, and fold change was determined relative to the control mRNA levels in freshly isolated TG-elicited macrophages (0). 0*, unstimulated macrophages adherent for 2 h before the addition of IL-4. Results shown are mean ± SD of three independent experiments. **(C)** Time-dependent expression of E-cadherin in fusing macrophages was quantified using immunofluorescence with anti-E-cadherin mAb followed by a secondary Alexa Fluor 546-conjugated goat anti-mouse antibody. Nuclei are stained with DAPI (teal). Representative images of E-cadherin expression for each time point are shown. The scale bar is 50 µm. **(D)** Quantification of fluorescence intensities of images of E-cadherin expression shown in **C**. Data shown are mean ± SD of three independent experiments. **(E)** Effect of EGTA on ZLS formation. Cells in the 5-day culture of IL-4–induced macrophages were treated with 1 mM EGTA for various periods (5–30 min). Vehicle buffer (0 min) was added to the control cells. *Upper panel*: After washing, samples were fixed and labeled with Alexa Fluor 546-conjugated phalloidin (white) and DAPI (teal). *Bottom panel*: High magnification views of the boxed areas shown in the upper panel. **(F)** The total lengths of ZLSs in the control (0 min) and EGTA-treated cells. Results shown are mean ± SD of three independent experiments. **(G)** Dose-dependent effect of EGTA on ZLS formation. Cells in the 5-day culture of IL-4–induced macrophages were treated with different concentrations of EGTA (0–1 mM) for 5 min, and the total lengths of ZLSs were determined. Results shown are mean ± SD of three independent experiments. \*\*\**p* < .001

To investigate the contribution of E-cadherin to the adjoining of the plasma membranes within the ZLS, we treated the cells in a 5-day culture with 1 mM EGTA. As shown in Figures 9E, the treatment of cells for 5 min resulted in almost complete dissociation of ZLSs. The effect of EGTA was dose-dependent with the concentration of EGTA as low as 0.06 mM reducing the formation of ZLSs by ∼90% (Figure 9G). These data indicate that E-cadherin is required for generating ZLSs and suggest that this junctional molecule is the chief mediator of ZLS formation.

## DISCUSSION

In this study, using an *in vivo* biomaterial implantation model we revealed previously unknown actin-based zipper-like structures that form at the sites of contact between large macrophage-derived MGCs. We reproduced the process of the formation of these structures *in vitro* and characterized their composition. Several lines of evidence indicate that ZLSs are transient dynamic structures that are formed from podosomes and disassembled into podosomes. First, the plasma membranes of two cells immediately preceding the ZLS are arrayed with podosomes. Second, podosome-specific proteins talin, vinculin, paxillin, cortactin, and integrin α_M_β_2_ are the components of ZLSs. Third, live-cell imaging using macrophages isolated from GFP- or mRFP-LifeAct mice enabled direct visualization of the origin of ZLS from podosomes. Lastly, the organization of ZLSs requires the Arp2/3 nucleation-promoting factors WASp and Cdc42, and the Arp2/3 complex, three proteins that are also involved in podosome formation. The intercellular space between the apposing plasma membranes within ZLSs is filled with the junctional proteins E-cadherin and nectin-2, suggesting that homophilic interactions between these molecules induce ZLS assembly.

In three dimensions, ZLSs show a characteristic pattern of actin distribution. Actin is mainly concentrated in large foci that are distributed with a regular spacing of ∼2.1 µm on each side of the plasma membrane of two abutting cells (Figure 3 and schematically shown in Figure 10). In addition, actin forms the second array consisting of small globules positioned in the immediate proximity to the plasma membrane. Video recordings revealed that large actin foci are assembled from podosomes that arrive and concentrate along the membrane and enlarge by recruiting adjacent single podosomes (Figure 5 and Video 1 and 2). As determined from the fluorescence micrographs of fixed specimens, the average diameter of large actin foci (∼1.5 µm in diameter) is greater than that of individual podosomes (0.7 ± 0.2 µm). This suggests that at the cell-substrate surface, large foci may contain 2–3 individual podosomes. However, since the height of large foci is ∼2.9 µm (Figure 3C) and the reported height of podosomes is 0.4–0.6 µm (Linder, 2007; Labernadie *et al*., 2010), the calculated volume of large actin globules and podosomes is ∼1.2 µm^3^ and ∼0.17 µm^3^, respectively, based on the assumption of their shape as a circular, slightly truncated cone (Linder, 2007). These values suggest that large actin globules can accumulate actin from ∼7–8 individual podosomes. Dynamic clusters of podosomes have previously been reported in LPS- and INFγ-stimulated IC-21 mouse macrophages in which podosomes fuse to form transient actin-containing structures that vary in size (Evans *et al*., 2003). Likewise, LPS/IFNγ treatment of human monocyte-derived macrophages causes podosomes to cluster (Poincloux *et al*., 2006). The size heterogeneity of the actin core has also been observed in dendritic cells, in which a population of podosomes undergoes fusion or fission (van den Dries *et al*., 2013a). Therefore, it appears that large actin foci in ZLSs arise by acquiring new podosomes. Furthermore, the relatively uniform dimensions of large actin foci suggest that only a limited number of podosomes can combine, indicating that their size is regulated.

**Figure 10.**
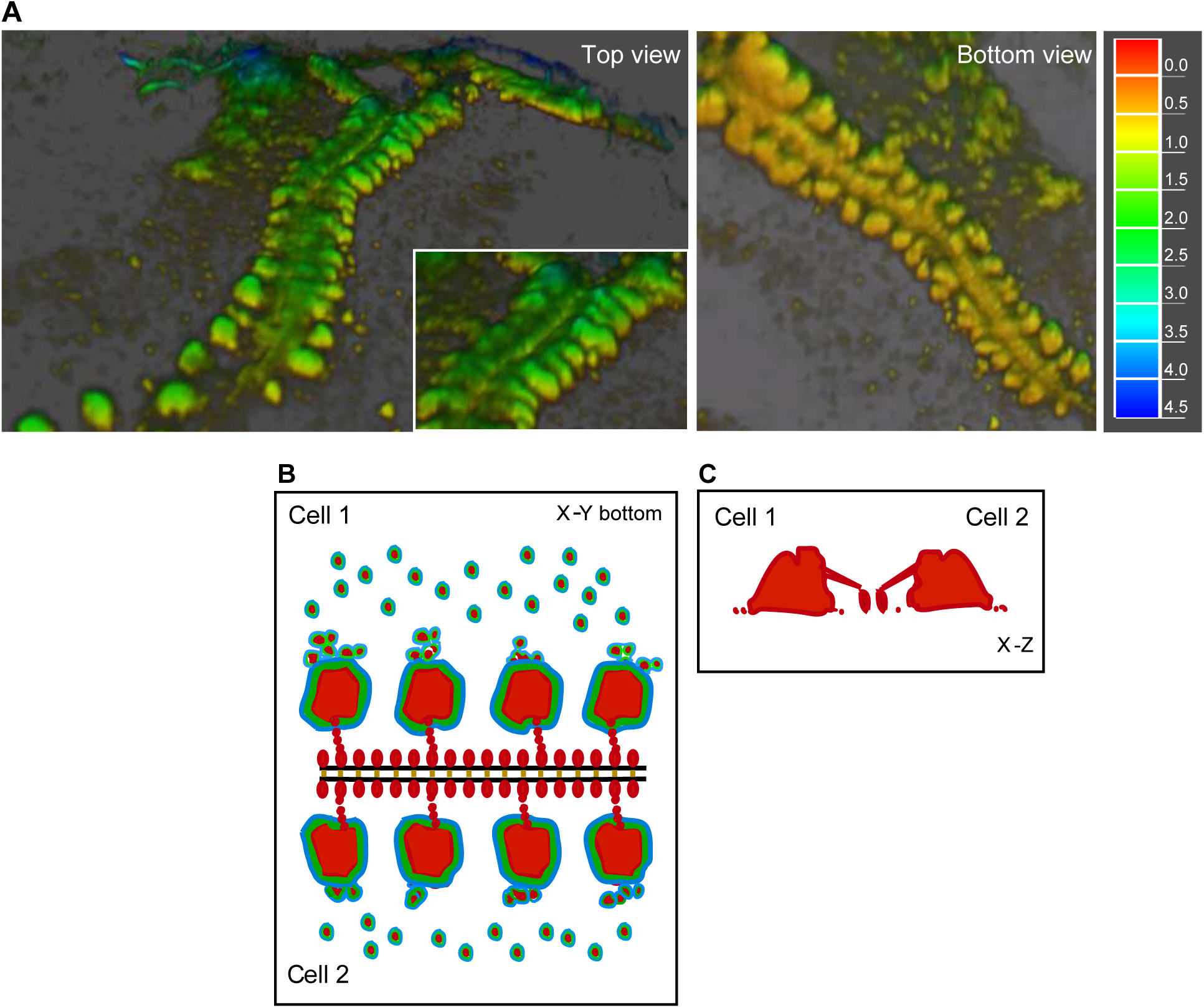
Schematic drawing of the organization of actin and associated proteins in a ZLS. **(A)** The 3D reconstruction of actin distribution in a ZLS formed by two abutting cells, based on confocal and structured illumination microscopy using cell labeling with Alexa-Fluor-conjugated phalloidin. *Left panel*: top view. *Right panel*: bottom view. The representation of data as a heat map is shown on the right (µm). (**B**) The distribution of actin (red) within the large and small membrane-proximal globules in a ZLS. Based on live-cell video microscopy, large globules form from a pool of individual podosomes present in the vicinity of the plasma membranes. The podosome proteins vinculin and talin (green and blue, respectively) surround the actin cores in large globules and in individual podosomes. E-cadherin (orange) holds the two plasma membranes (black lines). Afadin (not shown) and other unknown scaffold proteins may organize actin and its associated proteins into the highly symmetrical ZLS structure. (**C**) A vertical section across two large actin globules in a ZLS.

A striking feature of ZLSs is that actin foci, large or small, are correctly aligned against each other on both sides of the cell-cell interface and equidistant from the plasma membrane, implicating undefined scaffold proteins in ZLS formation. It is worth noting that a distinctive pattern of actin distribution is clearly seen only in the x-y plane and cannot be easily discerned in the z-direction, especially in the region adjacent to the plasma membrane (Figures 3 and 10). The z scans show that actin within large globules is connected with actin in small globules, although its nature, as well as that of small globules, is not clear. The humps also contain proteins typically found in podosomes, including talin, vinculin, paxillin, cortactin, and integrin α_M_β_2_ (Figures 6 and 10). The discrete position of vinculin and talin observed in the x-y plane is similar to that in podosomes; i.e. both proteins form rings surrounding actin cores in large foci (Figure 6, A-F). The distributions of cortactin and α_M_β_2_ differ from those of other proteins (Figure 6). Interestingly, both cortactin and α_M_β_2_ were observed in the midline, suggesting that these proteins are positioned close to the membrane. Furthermore, since α_M_β_2_ was detected using mAb M1/70 directed against the ligand-binding domain of integrin, these data indicate that α_M_β_2_ is present in the intercellular space. In the scheme shown in Figure 10, we take this distribution of actin and ZLS-associated vinculin and talin into account, although further studies are required to determine the complex organization of a ZLS.

ZLSs formed only after a prolonged period of adhesion of MGCs to the surface of biomaterials *in vivo* and after several days of incubation with IL-4 in a culture medium *in vitro*. By this time, the majority of MGCs appear as large spread cells that form extensive contacts with each other, apparently facilitating ZLS formation. Our live-cell imaging analyses revealed that ZLSs are dynamic structures with an average lifespan of ∼13 min. Notably, the assembly of ZLSs seems to occur in a sequential manner, visually resembling a fastener that moves in one direction, “zippering up” plasma membranes. The zippering mechanism is apparently enacted by E-cadherin and nectin-2, which are detected in the intercellular space of a ZLS. In agreement with previous data (Moreno *et al*., 2007; Van den Bossche *et al*., 2009), E-cadherin mRNA was poorly detected in freshly isolated macrophages but was gradually expressed and peaked within 12 h of IL-4 addition and remained stable after 5 days (Figure 9). Moreover, while the protein was weakly expressed on the surface of freshly plated macrophages, its expression was readily detectable by 24 h and gradually upregulated over 5 days (Figure 9). Since ZLSs were first detected at day 4, either the critical level of E-cadherin or the required level of cell spreading or both may be an essential requisite of ZLS formation. Similar to E-cadherin, nectin-2 mRNA was not detected in freshly isolated macrophages but expressed several hours after incubation in the presence of IL-4. Nonetheless, E-cadherin appears to play a dominant role in ZLS formation inasmuch as sequestration of Ca^2+^ by EGTA effectively disrupted ZLSs (Figure 9).

E-cadherin and nectin-2 are typical components of adherens junctions (AJ) in epithelial and other cells (Takai *et al*., 2008; Harris and Tepass, 2010; Takeichi, 2014). In AJs, Ca^2+^-dependent cadherin molecules associate with the actin cytoskeleton, strengthening intercellular adhesions. Moreover, another intercellular adhesion system consisting of the Ca^2+^-independent nectin molecules and nectin-binding adaptor protein afadin, which connects nectin to actin, plays a role in the organization of AJs, either cooperatively with or independently of cadherin (Takai *et al*., 2008). E-cadherin and nectin-2 have previously been identified in monocytes/macrophages (Pende *et al*., 2006; Moreno *et al*., 2007; Van den Bossche *et al*., 2009). The surprising finding of the present study is that these molecules, together with afadin, were found in ZLSs suggesting that they can form adherens junction-like structures in mature MGCs. Epithelial AJs have traditionally been classified into two major groups based on their dynamics and on how they associate with actin filaments (Franke, 2009; Takeichi, 2014). The linear adherens junctions (lAJ) that connect mature epithelial cells link with a bundle of linear actin filaments that runs parallel to the cell borders and are relatively stable. The punctate adherens junctions (pAJ) that are found at the edges of epithelial colonies and in other cell types, associate with radial actin bundles and are mobile, morphologically unstable structures (Takai *et al*., 2008; Takeichi, 2014) (Indra *et al*., 2018). E-cadherin is also observed at lateral cell-cell contacts below lAJ, where it associates with an amorphous actin network(Takeichi, 2014). There are several notable differences between the ZLS-type junctions in MGCs, and AJs in epithelial cells. First, in contrast to AJs, the transient nature of the ZLS-type junctions suggests that they are held by weak interactions. Although pAJs have been described as mobile structures, this behavior is referred to as the ability of cadherin molecules to be released and then reassemble within cadherin clusters (Indra *et al*., 2018). Hence, the dynamic behavior of pAJ is related to the continuous turnover of cadherin molecules while still maintaining their overall stability. In contrast, an entire ZLS can form and disassemble in one location while a new ZLS appears in another location. Second, the distribution of actin in ZLSs is distinct from that observed in AJs. Neither parallel actin filaments similar to those in lAJ nor pAJ-associated long filaments that perpendicularly terminate at the plasma membrane have been detected in ZLSs. Rather, actin is organized within regular foci that resemble large podosome clusters. Third, in contrast to lAJs that may encircle cells, ZLSs form discontinuous segments at the sites where MGCs contact each other. In this regard, ZLSs are reminiscent of cell-cell junctions formed in fibroblasts and other motile cells, in which they are observed as punctate and streak-like structures (Yonemura *et al*., 1995; Takeichi, 2014). Fourth, the inter-membrane space in a ZLS determined by TEM was 7.7 ± 1.3 µm, significantly narrower than 15–25 µm in epithelial AJs (Farquhar and Palade, 1963; Miyaguchi, 2000). Thus, despite the presence of some typical AJ proteins and actin, many features of the ZLS-type junctions in MGCs distinguish them from AJs and other cell-cell junctions, suggesting that these adhesive structures may represent a novel type of AJ. Recently, a report has shown that mononuclear macrophages in the granuloma of zebrafish infected with mycobacteria can form adherens junctions (Cronan *et al*., 2016). Since MGCs are the characteristic feature of granulomas in tuberculosis and E-cadherin expression has been detected in macrophages from human and mouse samples (Cronan *et al*., 2016), it will be interesting to examine whether MGCs in granulomas also form ZLSs.

In addition to E-cadherin and nectin-2, integrin α_M_β_2_, which is known to associate with podosomes in macrophages (Duong and Rodan, 2010; van den Dries *et al*., 2013a), was found in the intercellular space of ZLSs. Integrin α_M_β_2_ is a multi-ligand receptor that can engage several counter-receptors, including members of the ICAM and JAM protein families (Diamond *et al*., 1990; Santoso *et al*., 2002). It is possible that α_M_β_2_ may contribute to heterophilic interactions that hold two membranes in ZLSs together. We have recently demonstrated that α_M_β_2_ can interact with SIRPα (also known as the macrophage fusion receptor, MFR) (Podolnikova *et al*., 2016), which belongs to the Ig superfamily similarly as ICAMs and JAMs. However, while SIRPα was strongly expressed in ZLSs (Supplemental Figure 4A), it was not present in the midline, suggesting that SIRPα and α_M_β_2_ do not colocalize. At present, the counter-receptor of α_M_β_2_ and its contribution to ZLS formation remain to be determined.

We have recently demonstrated that macrophage fusion occurs in three overlapping steps *in vitro* (Faust *et al*., 2017). Several hours after IL-4 induction, a founder population of mononuclear macrophages initiates fusion with neighboring mononuclear macrophages. These early multinucleated cells then fuse with neighboring mononuclear macrophages and, finally, MGCs fuse with surrounding MGCs to form syncytia. Our current studies suggest that the formation of ZLSs is a late-stage event in multinucleation. Since similar structures form in late MGCs on the surfaces of implanted biomaterials, these data indicate that ZLSs are not *in vitro* artifacts. However, the role of these enigmatic structures is unclear. It is well-known that when motile cells establish cell-cell junctions they cease movement and proliferation, a phenomenon referred to as contact inhibition of cell movement and proliferation (Fisher and Yeh, 1967; Bell, 1978). Since peritoneal macrophages do not divide, it is unlikely that ZLSs function in blocking cell proliferation signals. Therefore, the role of ZLSs in suppressing migration of MGCs as well as mononuclear cells that occasionally form ZLSs with MGCs can be theoretically envisioned. Our studies demonstrate that MGCs remain viable after establishing ZLSs. This observation is consistent with a well-known fact that cells that establish cell-cell junctions and stop moving continue to survive. Another possibility is that ZLSs may be involved in proteolysis. Podosomes have been associated *in vitro* with ECM degradation in many cell types, including macrophages (Linder *et al*., 2011). This is achieved by the recruitment and localized release of MMPs as well as serine and cathepsin proteinases. Among MMPs, MT1-MMP has been shown to degrade ECM in primary macrophages (Wiesner *et al*., 2010; Wiesner *et al*., 2014). Although we observed MT1-MMP in mononuclear macrophages and early MGCs, we were not able to detect this protease in ZLSs (Supplemental Figure 2C). At present, the involvement of other proteases that can be recruited with podosomes to ZLSs is unknown.

Recent studies have demonstrated ZLSs in the cultures of the murine macrophage cell line RAW267.4 undergoing osteoclastogenesis in the presence of RANKL and in osteoclasts induced in mouse bone marrow cells by RANKL/M-CSF (Takito *et al*., 2012; Takito *et al*., 2017). Although osteoclast ZLSs and the ZLSs observed in the current study share similar features, such as overall visual appearance, transient nature, and association with podosomal proteins, there are clear differences between these two structures. First, two types of ZLSs have been found in osteoclasts. The first type has been observed at the ventral membrane of a single multinucleated osteoclast that formed as a result of fusion of mononuclear cells and was interpreted as the structure joining actin rings of individual cells remaining after fusion. Such compartmentalization of the ventral membrane was not observed in IL-4–induced MGCs. We invariably observed ZLSs only between the plasma membranes of two large apposing MGCs (and rarely between an MGC and mono/binuclear cell). The ZLS structures in our experiments seem to resemble another type of ZLS that was observed in mature osteoclasts (Takito *et al*., 2017). Second, in contrast to MGCs, the ZLSs in osteoclasts were negative for E-cadherin, β-catenin and nectin-2 staining (Takito *et al*., 2012). Third, the dimensional parameters of ZLSs in mature osteoclasts are different from those in MGCs, with the average width of the ZLSs in osteoclasts being almost twice larger than that in MGCs (∼8.4 µm vs. ∼4.8 µm, respectively). Fourth, perhaps the most striking difference between the ZLSs in MGCs and those in osteoclasts is their mechanisms of formation. Although both structures are highly dynamic, the ZLSs in MGCs are formed from podosomes and disassemble into podosomes (Figure 4), whereas the ZLSs in osteoclasts result from continuous retrograde actin flow and are independent of the dissolution and reformation of podosomes (Takito *et al*., 2017). It has been proposed that this actin flow generates forces that push the plasma membranes of neighboring osteoclasts at cell-cell contact sites to generate the ZLSs. In contrast, our results suggest the ZLSs in MGCs are formed by the interaction of E-cadherin molecules through the zippering mechanism. Finally, the ZLSs in osteoclasts have been proposed to be involved in cell-cell fusion (Takito *et al*., 2012). This scenario differs from our findings. Since the ZLSs in MGCs materialize only in extended IL-4–induced cultures by the time the cell fusion largely ceases (Figure 2, G and H), it is unlikely that MGC ZLSs are involved in the fusion. Indeed, among the twenty ZLSs analyzed by live-cell imaging, we were able to detect only one fusion event that occurred at the site of the ZLS. Thus, the kinetic and morphological differences between osteoclast and MGC ZLSs indicate that the definitive description of ZLSs during osteoclastogenesis and MGC formation and their functional roles may require further analysis.

In conclusion, our *in vivo* and *in vitro* studies demonstrated the formation of highly ordered actin-based zipper-like structures that originated from podosomes and linked the plasma membranes of two large MGCs. Given the fact that macrophage-derived MGCs and their secreted products may modulate the foreign body reaction to implanted biomaterials and ultimately wound healing (Anderson *et al*., 2008) (Jones *et al*., 2008), the mechanisms underlying the assembly of podosome-derived ZLSs and their role in the foreign body reaction merit further analyses.

## MATERIALS AND METHODS

### Reagents

The rat mAb M1/70, which recognizes the mouse α_M_ integrin subunit, was purified from the conditioned media of hybridoma cells (obtained from The American Tissue Culture Collection, Manassas, VA) using protein A agarose. The mouse anti-talin (T3287) and anti-vinculin (V9131) mAbs and rabbit anti-I/S-afadin polyclonal antibody (A0224) were from Sigma (St. Louis, MO). The rabbit anti-paxillin (32084) and anti-JAM-A (180821) polyclonal antibodies and rabbit anti-cortactin mAb conjugated to Alexa Fluor 555 were from Abcam (Cambridge, MA). The rat anti-nectin-2 (502-57) mAb was from Santa Cruz Biotechnology (Dallas, TX). The mouse anti-E-cadherin mAb (24E10) and rabbit anti-Myosin IIa polyclonal antibody (34035) were from Cell Signaling (Danvers, MA). The rat anti-SIRPα/CD172a polyclonal antibody (P84; 552371) was from BD Bioscience (San Jose, CA). The rabbit anti-MT1-MMP-14 polyclonal antibody (14552-1-AP) was from Proteintech (Rosemont, IL). The rabbit anti-ZO-1 polyclonal antibody (61-7300) and secondary antibodies Alexa Fluor 488-conjugated goat anti-rabbit IgG and Alexa Fluor 633-conjugated goat anti-rat IgG were from Invitrogen (Carlsbad, CA). Vibrant DiD membrane-labeling reagent was from Thermo Fisher (Waltham, MA). Brewer’s thioglycollate (TG), wiskostatin, and the Arp2/3 inhibitors CK-548 and CK-636 were from Sigma (St. Louis, MO). IL-4 was from Genscript (Piscataway, NJ).

### Mice

C57BL/6J and WASp^-/-^ (B6.129S6-Was^tm1Sbs^/J) mice were purchased from The Jackson Laboratory (Bar Harbor, MA). EGFP- and mRFP-LifeAct mice (Riedl et al., 2010) were gifts from Dr. Janice Burkhardt and were used with permission from Dr. Roland Wedlich-Söldner. Myeloid cell-specific Cdc42^-/-^ mice were generated by crossing Cdc42^loxP/loxP^ mice with LysMcre mice, followed by screening the progeny for Cdc42 excision in myeloid leukocytes as previously described (Faust *et al*., 2019). All animals were given *ad libitum* access to food and water and maintained at 22 °C on a 12-hour light/dark cycle. Experiments were performed according to animal protocols approved by the Institutional Animal Care and Use Committees at Arizona State University.

### Biomaterial implantation

Segments (1.5 × 0.5 cm) of sterile polychlorotrifluoroethylene (PCTFE) were implanted into the peritoneum of age- and sex-matched mice. Animals were humanely sacrificed 14 days later, and explants were analyzed for the presence of MGC as previously described (Faust *et al*., 2019). Prior to explantation, 2 ml of PBS containing 5 mM EDTA was aseptically injected into the peritoneum and cells in the peritoneum were collected by lavage. The number of cells in the peritoneum at the time of explantation was determined by counting with a Neubauer hemocytometer. Experiments were conducted in triplicate on three independent days.

### Macrophage isolation

Macrophages were isolated from 8–12-week-old male and female age- and sex-matched mice injected (I.P.) with 0.5 mL of a sterile 4% Brewer’s thioglycollate (TG) solution. All animals were humanely sacrificed 72 h later, and macrophages were isolated by lavage with ice-cold phosphate-buffered saline (PBS, pH 7.4) containing 5 mM EDTA. Macrophages were counted with a hemocytometer immediately thereafter.

### IL-4–induced macrophage fusion

Macrophage fusion was induced as previously described (Faust *et al*., 2019). Briefly, peritoneal cells (5 × 10^6^ cells/ml) in Hank’s Balanced Salt Solution (HBSS; Cellgro, Manassas, VA) supplemented with 0.1% bovine serum albumin (BSA) were applied to acid-cleaned or paraffin-coated glass coverslips [prepared as described by Faust *et al*. (2017 and 2018)]. Cells were incubated in 5% CO_2_ at 37 °C for 30 min. Non-adherent cells were removed by washing the culture three times with HBSS, and then the adherent cells were cultured with DMEM/F12 containing 15 mM HEPES (Cellgro, Manassas, VA), 10% FBS (Atlanta Biological, Flowery Branch, GA) and 1% antibiotics (Cellgro, Manassas, VA). After 2 h, 10 ng/ml IL-4 was added to the cultures until the indicated time points. For the 5-day cultures, media were changed on day 3. The fusion indices were determined from the images of cells labeled with Alexa Fluor 568-conjugated phalloidin and DAPI as previously described (Faust *et al*., 2019). The fusion index is defined as the fraction of nuclei within MGCs and expressed as the percentage out of the total nuclei counted. A total of 18–20 images (40×) that contained approximately 100–200 cells were analyzed for each experimental condition. The lengths of ZLSs were determined with ImageJ software (National Institutes of Health). Photomicrographs of representative fields were obtained with a Leica SP8 microscope (Leica Microsystems Inc. Buffalo Grove, IL).

### Phase-Contrast Video-Microscopy

Wild-type, WASp-deficient, and Cdc42-deficient macrophages (5 × 10^6^/ml) isolated from the peritoneum of mice 3 days after TG injection were plated on the surface of paraffin-coated coverslips or a section of polychlorotrifluoroethylene (PCTFE), and cell fusion was induced by adding 10 ng/ml IL-4. Dishes were transferred from the cell culture incubator to a stage-top incubator calibrated to maintain a humidified atmosphere of 5% CO_2_ in air at 37 °C. Phase-contrast images were collected with a 20× objective every 30 s using an EVOS FL Auto (Thermo Scientific, Waltham, MA) and transferred to ImageJ to generate movies.

### Live-cell fluorescence microscopy

EGFP- or mRFP-LifeAct macrophages were isolated from the peritoneum of TG-induced mice. Macrophages (10^6^/0.5 ml) were plated on the surface of coverslips adsorbed with paraffin, and incubated in the presence of 10 ng/ml IL-4. After 3 days, half of the DMEM/F12 medium was replaced with fresh medium without IL-4, and the cell culture was incubated for an additional 2 days. The dish was placed into a live-cell imaging chamber supplied with 5% CO_2_ at 37 °C, and live-imaging was conducted for 6 h. Images were acquired by Leica SP8 using a 40×/1.3 NA oil objective every minute using a HyD hybrid detector. The acquired images were processed and converted into movies using ImageJ software.

### Immunofluorescence

At the indicated time points, cells cultured on clean glass or paraffin-adsorbed coverslips were fixed with 2% paraformaldehyde in PBS for 30 minutes, permeabilized with 0.2% Tween-20 in PBS for 15 min at 22 °C, and then washed with PBS. The permeabilization step was omitted for the staining of transmembrane proteins. The cells were incubated overnight at 4 °C with the primary antibodies using the dilutions recommended by the manufacturers. Incubations with Alexa Fluor (488, 568, or 647)-conjugated secondary antibodies were performed at room temperature for 4 h. Cells were also stained with 15 nM Alexa Fluor 568-conjugated phalloidin (Thermo Scientific, Waltham, MA) for 30 min at 22 °C to detect F-actin. Cells were washed twice with PBS and incubated with DAPI. ProLong Diamond Antifade Mountant was used to mount the cells on a glass slide (Thermo Scientific, Waltham, MA). Images were acquired by using Leica SP8 and Zeiss LSM800 confocal microscopes (Carl Zeiss Vision Inc., San Diego, CA) with 40×/1.3 NA and 60x/1.4 oil immersion objectives, respectively. Super-resolution images were acquired using a Nikon SIM (Nikon Instruments Inc., Melville, NY) with an SR Apo TIRF 100x/1.49 NA oil immersion objective.

### RT-PCR analysis of *E-cadherin* and *Nectin-2* expression

Macrophages were induced to fuse by IL-4 for the indicated periods. Total RNA was extracted using TRIzol reagent (Invitrogen) and resuspended in 20 μl of RNase-free water supplemented with 0.1 mM EDTA. Afterward, 1 μg of total RNA was used to construct cDNA using SuperScript III Reverse Transcriptase (Invitrogen). PCR was performed with the generated cDNA and premixed 2× Taq polymerase solution (Promega) in an MJ Mini Thermal Cycler (BioRad). The levels of target mRNAs were normalized by *Tuba1b* mRNA level. The primer sets for PCR analyses were purchased from Integrated DNA Technologies (Iowa, USA) and included those for *Tuba1b,* 5’-CAGGTCTCCAGGGCTTCTTG-3’ (*forward*) and 5’-GAAGCATCAGTGCCTGCAAC-3’ (*reverse*); for *Cdh1* 5’-CGGGACTCCAGTCATAGGGA-3’ (*forward*) and 5’-ACTGCTGGTCAGGATCGTTG-3’ (*reverse*); and for *Nectin2* 5’-GTTCAGCAAGGACCGTCTGTC-3’ (*forward*) and 5’-ATCGTAGGATCCTCTGTCGC-3’ (*reverse*). The semi-quantitative digital analysis was performed using ImageJ 1.52n software. Pixel density for each band was calculated in arbitrary units and expressed as a fold change relative to the target mRNA level in macrophages in suspension (denoted 0).

### TEM

TG-elicited peritoneal macrophages were seeded on a PCTFE section (Welch Fluorocarbon, Dover, NH) and cultured in DMEM/F12 supplemented with 15 mM HEPES, 10% FBS, and 1% antibiotics. After 2 h, 10 ng/ml IL-4 was added to cultures, and cells were incubated for 5 days. Cells were fixed with 2.5% glutaraldehyde in 0.1 M PBS (pH 7.4) at 4 ^°^C overnight and then treated with 1% OsO_4_ in 0.1 M PBS for 1 h. Subsequently, cells were washed with 0.1 M PBS and then dehydrated using acetone. Finally, cells were flat-embedded into Spur’s EPOXY Resin, and 70-nm sections were obtained by slicing parallel to the site of the cell attachment to substrate. Sections were post-stained with uranyl acetate and Sato’s lead citrate. Micrographs were taken using a Philips CM 12 TEM with a Gatan model 791 camera.

### Statistical Analyses

Unless otherwise indicated, results are shown as mean ± SD of three independent experiments. Multiple comparisons were made by using ANOVA followed by Tukey’s or Dunn’s post-test using GraphPad Instat software. Where applicable, means were compared with each other by using Student’s *t*-test. Data were considered significantly different if *p* < .05.

## Acknowledgment

We thank James Faust for helpful advice on performing phase-contrast live-cell video experiments and Page Baluch for anti-ZO-1 antibodies. We acknowledge the use of facilities within the Eyring Materials Center at Arizona State University supported in part by NNCI-ECCS-1542160. Image data were collected using a Leica TCS SP5 LSCM (the National Institutes of Health SIG award S10 RR027154) and Leica TCS SP8 LSCM (the NIH SIG award S10 OD023691) housed in the W.M. Keck Bioimaging Facility at Arizona State University.

## SUPPLEMENTAL FIGURE LEGENDS

**Supplemental Table 1.**
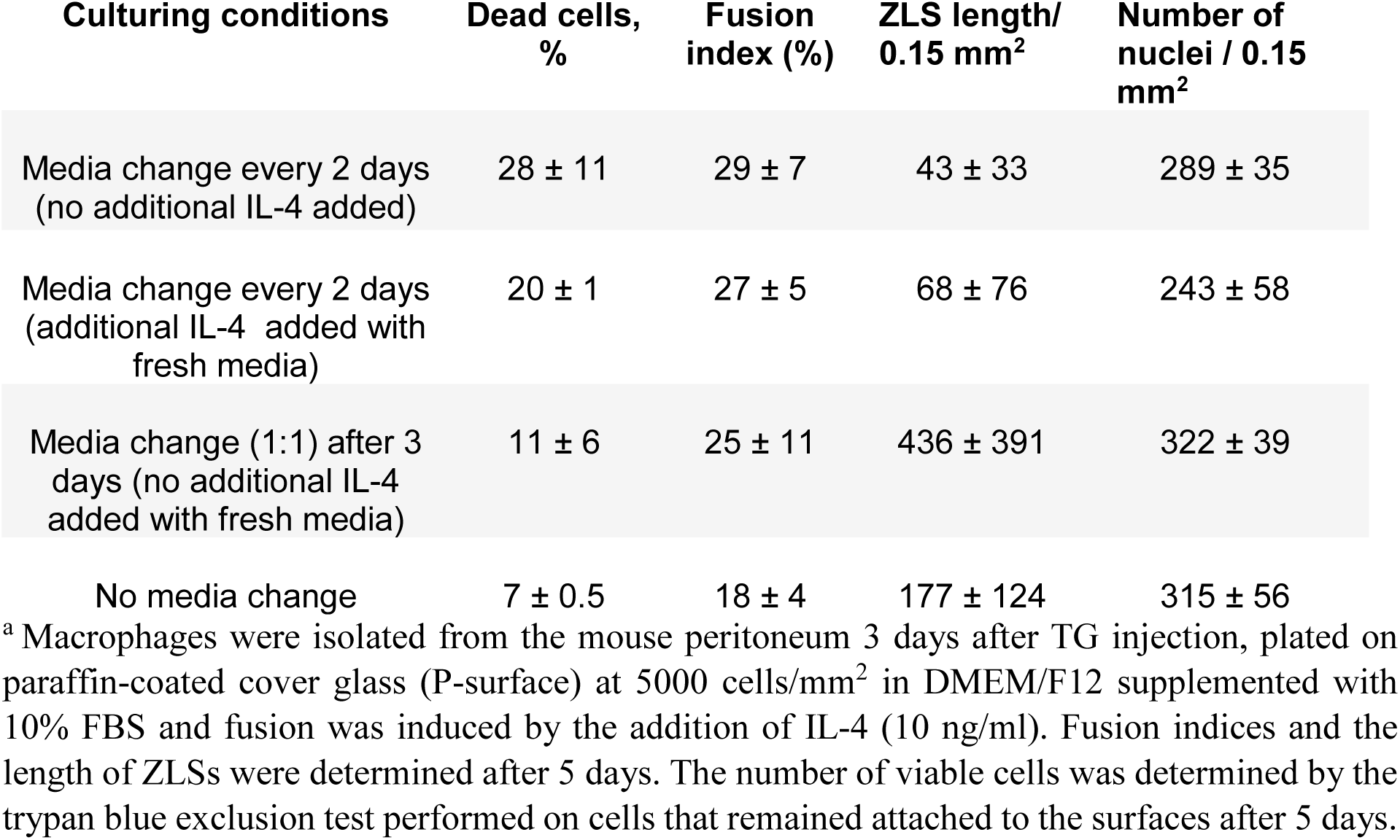
Viability of macrophages cultured under different conditions^a^

**Figure 1.**
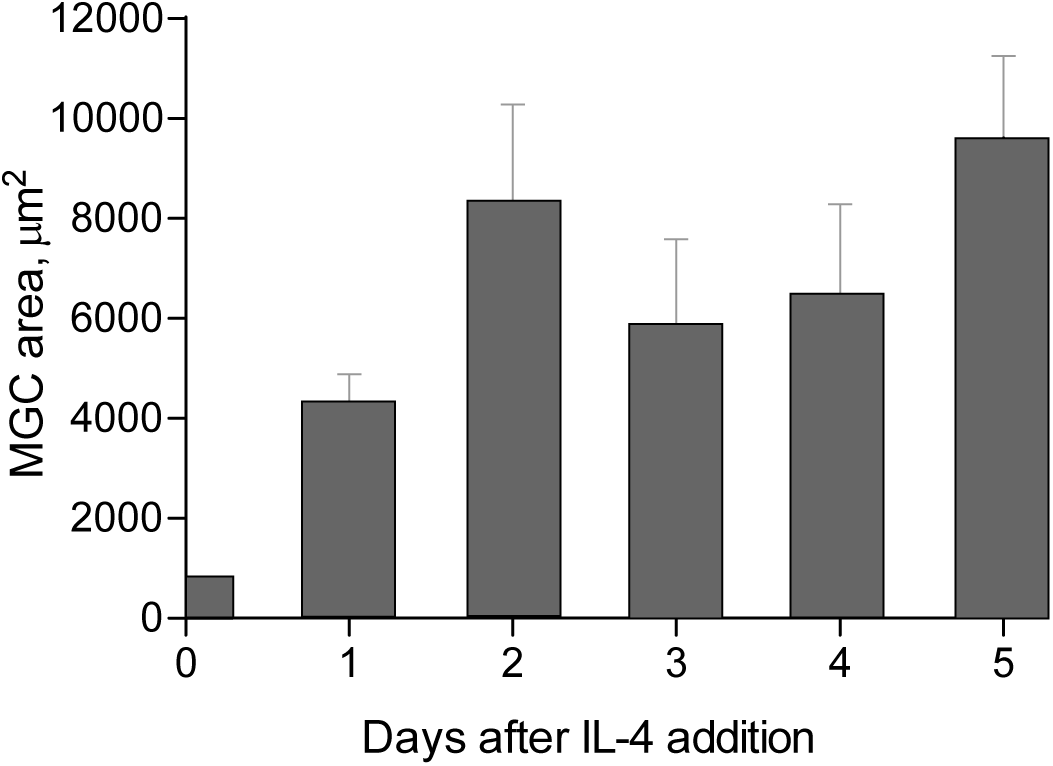
Determination of the MGC surface area. TG-elicited peritoneal macrophages were plated on P-surface. After the addition of IL-4, the cells were fixed at different time points and incubated with Alexa 488-conjugated phalloidin and DAPI. The samples were imaged by confocal microscopy. Results shown are mean ± SD of three independent experiments. Three-to-five random 20× fields were used per sample to measure the MGC area using ImageJ software.

**Figure 2.**
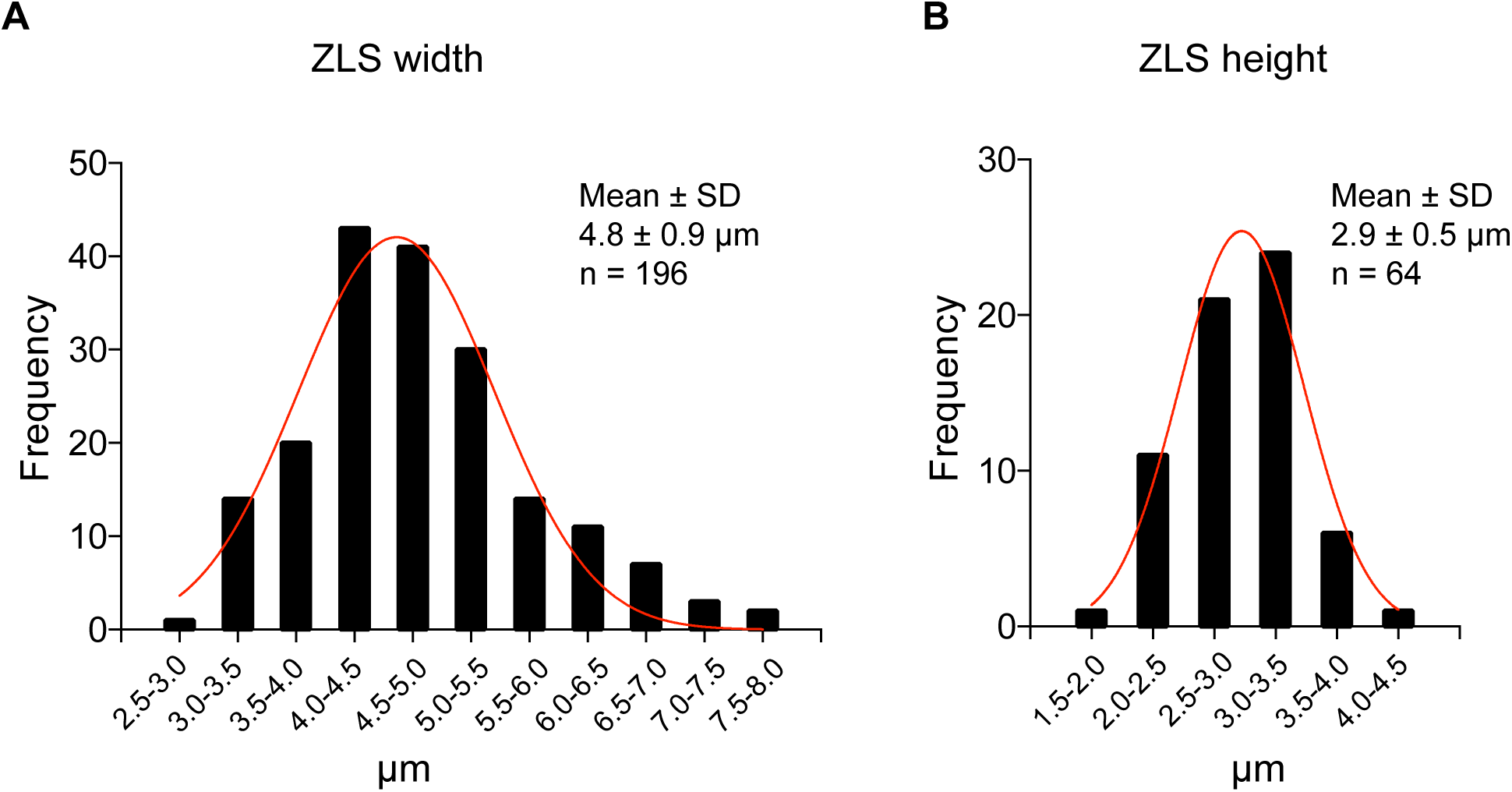
Dimensional parameters of ZLSs. The histograms showing the normal distribution of ZLS width (A) and height (B) were generated based on confocal images and produced using GraphPad Prism Software.

**Figure 3.**
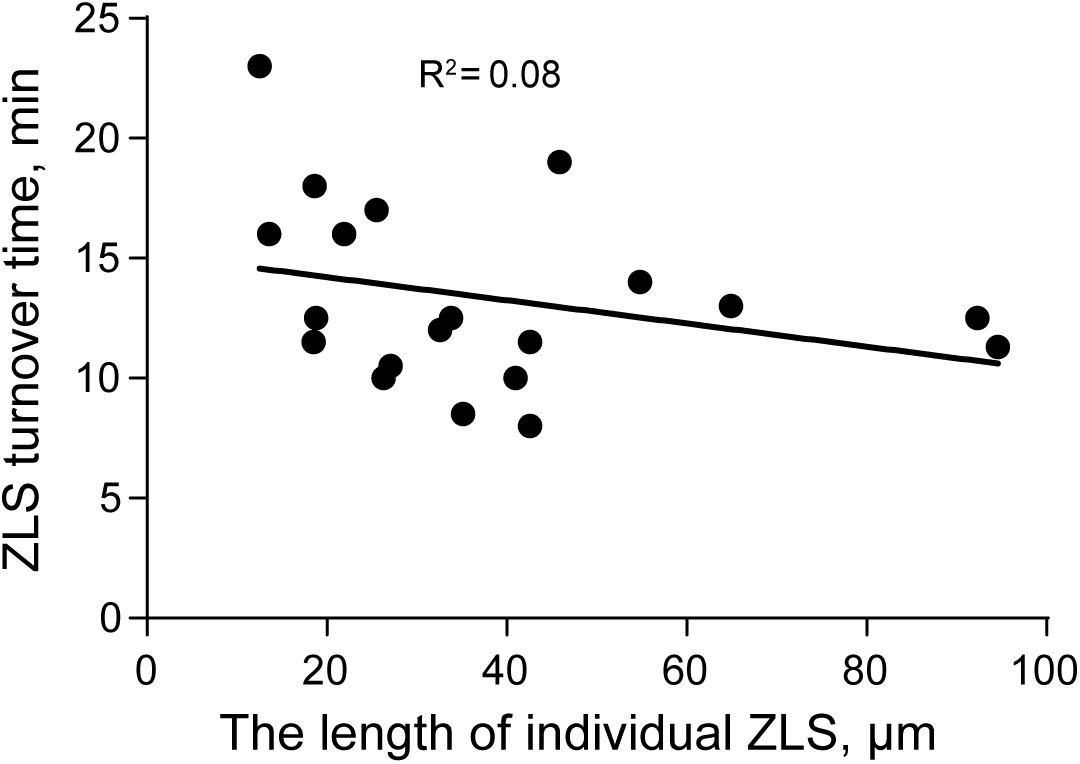
Correlation between the ZLS turnover time and ZLS length. Live-cell video microscopy images were analyzed to determine the correlation between the ZLS turnover time (the time required for the full assembly and disassembly of a ZLS) and the ZLS length (µm). No significant correlation was found.

**Figure 4.**
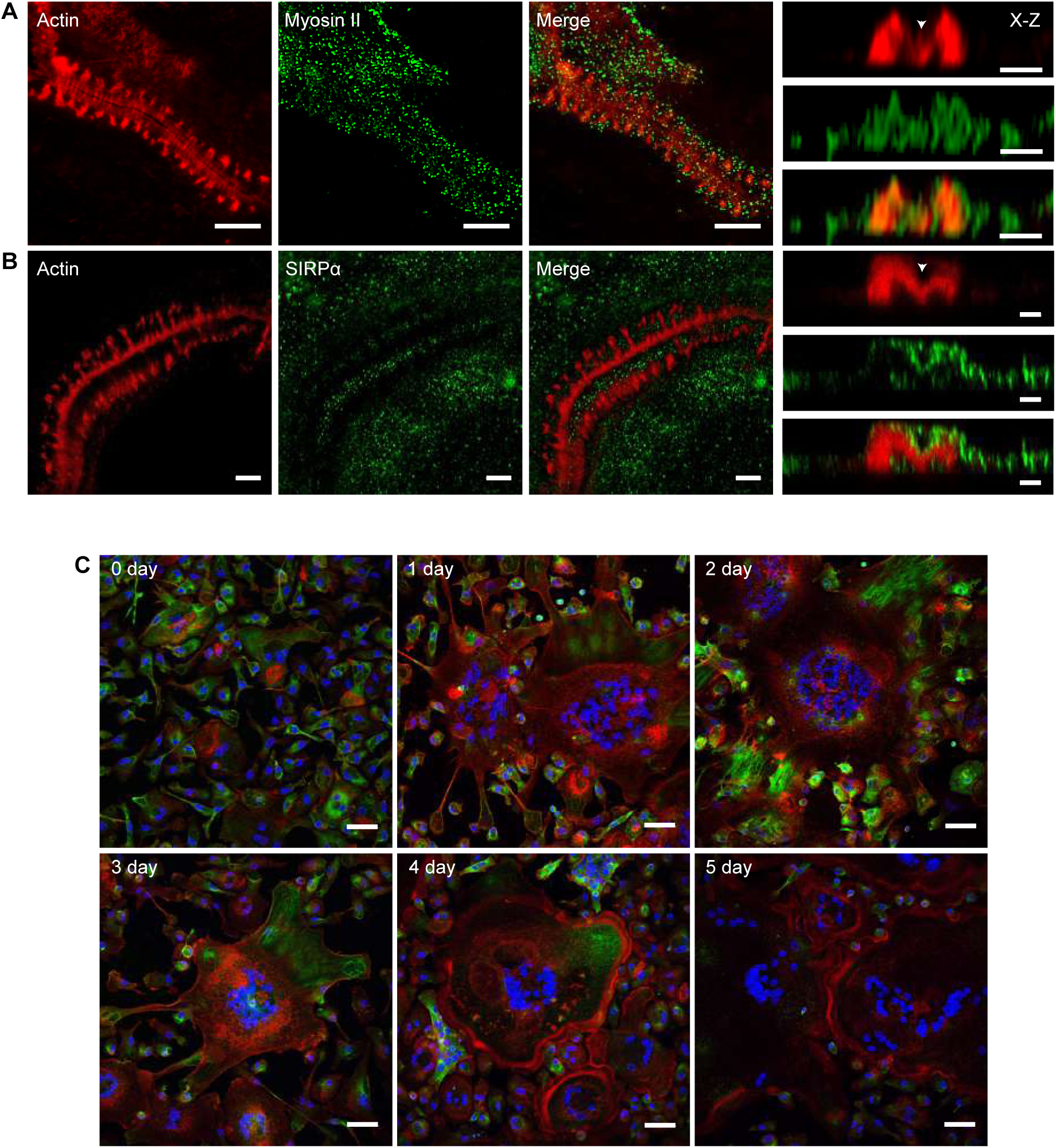
Distribution of Myosin II (A) and SIRPα (B) within ZLS structures. Representative confocal images of MGCs in the 5-day culture induced by the addition of IL-4 and incubated with anti-Myosin II **(A)** or SIRPα **(B)** mAbs. The x-z projections are shown in the right panels. The scale bars for x-y and x-z projections are 5 µm and 2.5 µm, respectively. The images were acquired using a 63× oil objective. **(C)** The kinetics of MT1-MMP-14 distribution in mononuclear macrophages and MGCs. TG-induced peritoneal macrophages were cultured in the presence of IL-4 for 1-5 days, fixed and incubated with antibodies against MT1-MMP-14. The images were obtained by confocal microscopy using a 40× oil objective. The scale bar is 50 µm.

**Figure 5.**
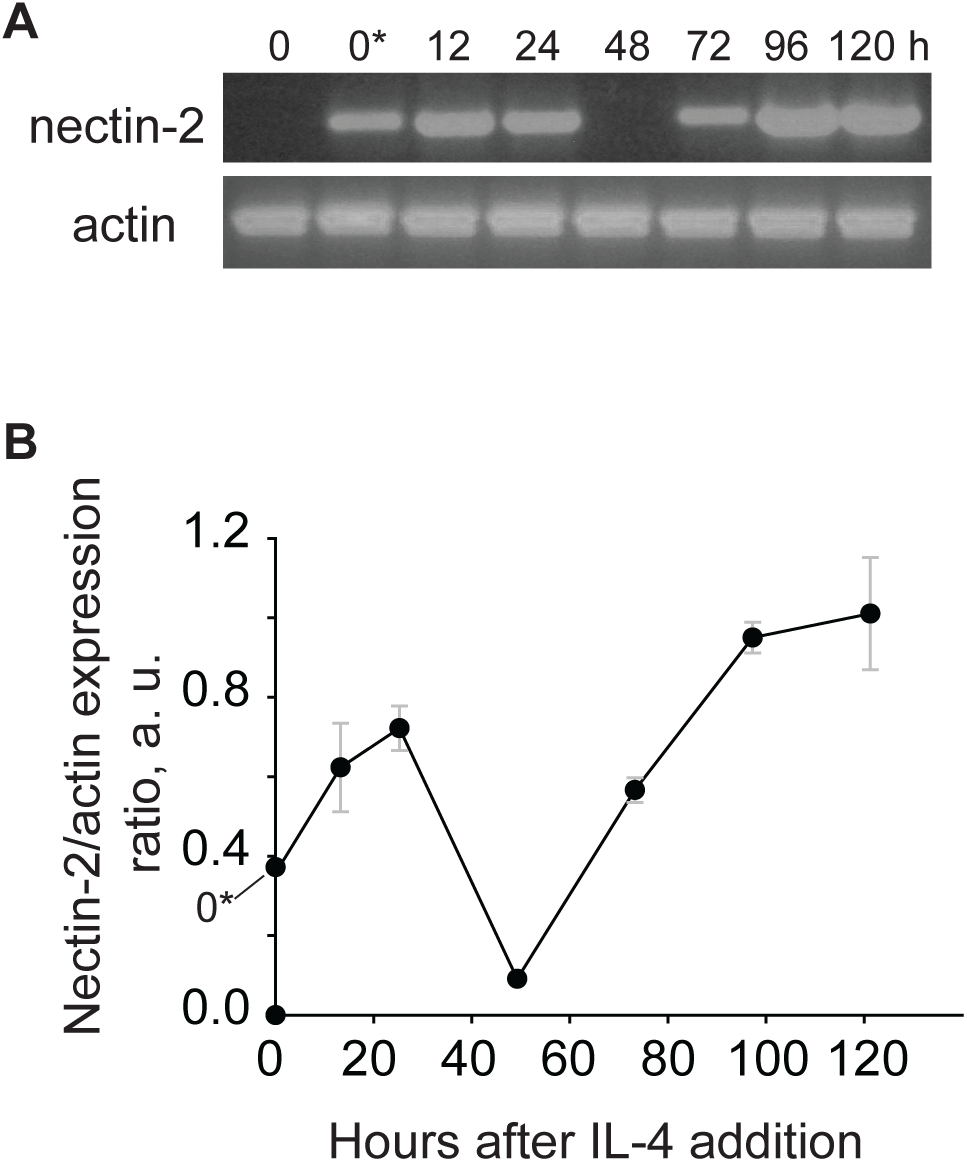
The kinetics of nectin-2 mRNA expression in fusing macrophages. **(A, B)** Time course of E-cadherin mRNA expression in macrophages undergoing fusion in the presence of IL-4, as determined by RT-PCR. Signal intensities were normalized to that of α-actin, and fold change was determined relative to the control mRNA levels in freshly isolated TG-elicited macrophages before adhesion (0). 0* denotes unstimulated macrophages adherent for 2 h before the addition of IL-4. Results shown are mean ± SD of two independent experiments.

## Notes

Support: This work was supported by the NIH grants HL63199 (TU) and R00NS076661 and R01NS097537 (JN)

